# Tissue-scale mechanics controls differentiation strategy and dynamics of epithelial multilayering

**DOI:** 10.64898/2026.02.08.704529

**Authors:** Clémentine Villeneuve, Somiealo Azote Epse Hassikpezi, Marga Albu, Matthias Rübsam, Leah C. Biggs, Sabrina Vinzens, Kai Kruse, Anubhav Prakash, Peter Zentis, Elizabeth Lawson- Keister, Gautier Follain, Johanna Ivaska, Carien M. Niessen, M. Lisa Manning, Sara A. Wickström

## Abstract

Generating and maintaining multilayered epithelia requires coordinated cell division, differentiation, and tissue architecture, yet the precise mechanisms of multilayering remain unclear. Using the developing mouse epidermis, we show that basal stem cells adopt distinct multilayering strategies depending on tissue mechanics. Combining quantitative morphometry, embryo live imaging and physical modeling, we observe that early in development, the epidermis is fluid-like, allowing undifferentiated cells to move suprabasally through perpendicular divisions or basal detachment before differentiating. As the tissue matures and rigidifies, a mechanical barrier is established that only allows upward movement of basal cells that have committed to differentiation. The final step of this commitment is delamination that requires Notch signaling, triggered by increased tissue stiffness and jamming. This mechanical regulation orchestrates a feedback loop that induces cell upward motion precisely when the basal layer becomes crowded. Together, our findings identify tissue mechanics as the key determinant of how tissues drive multilayering and reveal mechanically regulated Notch signaling as a driver of epidermal delamination.

## Introduction

Stratified epithelia such as the skin epidermis are multilayered tissues where the basal layer consists of undifferentiated epithelial stem cells adhering to the basement membrane, while the suprabasal layers contain post-mitotic, differentiated cells ^1–3^. Unlike other dynamically self-renewing epithelia such as the intestine, in which clear tissue-scale geometric cues such as curvature compartmentalize the stem cell away from differentiated progeny ^4^, it remains unclear how stratified epithelia establish and organize this spatial organization and differentiation dynamics during development and into adulthood.

To initiate or maintain epidermal stratification, the basal stem cells generate suprabasal cells via two distinct strategies: perpendicular divisions or detachment from the basal layer, termed delamination ^2,3,5^. Early in development, proliferation rates in the epidermis are high - to match the rapid growth of the organism it covers, and to generate sufficient suprabasal cells for efficient establishment of a functional life-essential barrier ^6^. During this period, perpendicular cell divisions generate suprabasal daughter cells ^7,8^, while also some delaminations are observed ^9,10^. In contrast, during adult homeostasis differentiation does not depend on cell divisions, with cells moving suprabasally exclusively through delamination to replenish the most superficial, desquamating cornified layers, followed by parallel divisions to maintain basal cell density ^11,12^. Key open questions remain: Why does this switch in strategies occur? And how is upward movement robustly regulated in the absence of obvious niche inputs like tissue curvature?

Here we demonstrate that multilayering strategy is controlled by a mechanical barrier that is established during development and hinders basal cells from moving upward. We establish a new approach to quantitively measure this mechanical barrier by combining a 3D biophysical model for cell and tissue mechanics with imaging of cell and tissue geometries in the basal layer and at the basal-suprabasal interface. We find that at early developmental stages, embryonic (E) day 13.5/14.5, there is little mechanical distinction between basal and suprabasal layers allowing basal cells to easily place cells suprabasally either through perpendicular division or through rapid detachment assisted by fluctuations from cell divisions. Around E15.5, a mechanical boundary between the basal and suprabasal layers emerges, and basal cells must now undergo profound transcriptional reprogramming– e.g., commit to delamination – in order to cross this boundary. How is this decision to commit regulated? Our data suggests an elegant feedback mechanism involving Notch signaling.

Tissue crowding and stiffening trigger a transient Notch signal in a small subset of cells, causing these cells to execute a transcriptional switch to complete delamination. Collectively, our study shows that changes in tissue mechanics that drive intercompartment boundary formation are essential determinants for a switch from early embryonic to a mature differentiation and multilayering strategy. Moreover, we identify Notch signaling as an integrative feedback relay to precisely time the transition between the mechanically separated basal and suprabasal compartments within the mature stratified epithelium.

## Results

### Basal cells employ distinct embryonic stage-specific strategies to generate and compartmentalize suprabasal layers

To understand which mechanisms are used for epidermal multilayering, we employed a whole embryo live imaging system ^9,13^ to visualize long term (over 8 h) cell dynamics at E14.5 and E15.5 (Fig. 1A). While perpendicular divisions were prevalent at E14.5 as previously reported ^8^, parallel oriented cell divisions became predominant at E15.5 with the overall frequency of cell divisions decreasing (Fig. 1B). As reported ^6^, cell divisions also occurred suprabasally at E14.5 but much less at E15.5 (Fig. 1B, Supplementary Fig. 1A). Imaging of fixed tissues showed that significantly more undifferentiated cells were found in the first suprabasal layer at E14.5 than at E15.5 (Supplementary Fig. 1B), as detected by the absence of the differentiation marker Keratin (K)10 ^14^ in suprabasal cells.

**Figure 1.**
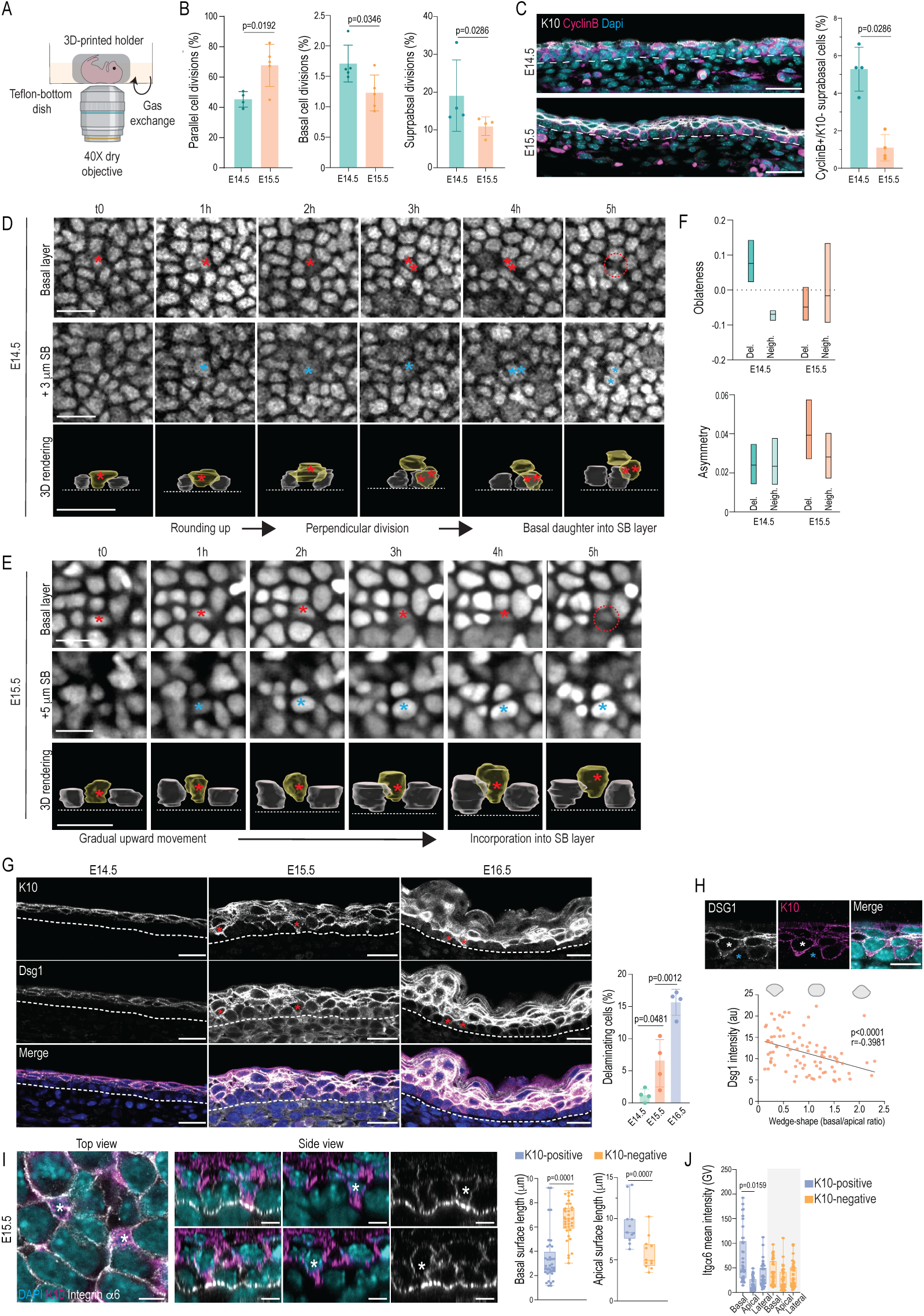
Basal cells employ distinct embryonic stage-specific strategies to generate suprabasal layers. **A.** Schematics of the *ex vivo* embryo imaging setup. **B.** Quantification of parallel (left panel), basal (middle panel) and suprabasal (right panel) cell divisions in E14.5 and E15.5 embryos (mean ±SD; n=4 (E14.5) /5 (E15.5) embryos; Student’s t-test (left and middle panel), Mann-Whitney (right panel)). **C.** Representative images and quantification of CyclinB-positive undifferentiated cells (K10- neg) in the suprabasal layers of E14.5 and E15.5 embryos (30 µm; mean ±SD; n=4 embryos/stage; Mann-Whitney). **D.** Representative snapshots and 3D rendering (lower panel) from live imaging movies of E14.5 epidermis from nTomato reporter mice. Red asterisk marks cell departing from basal layer; double red asterisk marks daughter of perpendicular division that subsequently moves suprabasally, blue asterisks marks appearance of cells in the suprabasal layer, red dotted circle marks completed delamination. Images representative of 3 embryos, scale bars 20 µm. **E.** Representative snapshots and 3D rendering (lower panel) from live imaging movies of E15.5 epidermis from nTomato reporter mice. Red asterisk marks cell departing from basal layer; blue asterisk marks appearance of this cell in the suprabasal layer, red dotted circle marks completed delamination. Images representative of 3 embryos, scale bars 20 µm. **F.** Quantification of nuclear oblateness (roundness; upper panel) and asymmetry of deformation (wedge-shape; lower panel) of delaminating cells (Del.) and their immediate neighbors (Neigh.) from 3D renderings. Note that while at E14.5 delaminating cells round up, at E15.5 they elongate in an asymmetric manner (min-to-max box plots; n=3 embryos/stage). **G.** Representative images and quantification of tissue sections from E14.5 - 16.5 embryos stained against K10, Dsg1 and DAPI. Note emergence of K10-Dsg1- positive cells at 15.5 (red asterisks). Dotted line marks basement membrane (scale bars 20 µm; mean ±SD; n=4 embryos; Two-way ANOVA/Tukey’s). **H.** Representative images and quantification of tissue sections from E15.5 embryos stained against K10, Dsg1 and DAPI. Note correlation of wedge-like shape and apical Dsg1 intensity. White asterisks mark wedge-like cells with high apical Dsg1, blue asterisks mark cuboidal neighboring cell (scale bars 10 µm; mean ±SD; n=87 cells pooled across 3 embryos; Simple linear regression; Pearson’s correlation). **I, J.** Representative images and quantification of delaminating cells in E15.5 embryo 3D whole mounts stained against integrin α6 (Itga6), K10 and DAPI. Note wedge-like shape and integrin α6-positive basal footprint of the delaminating cell (white asterisks; scale bars 5 µm; min-max box and whiskers; n=37 cells pooled across 3 embryos; Mann- Whitney (left, middle panel) Friedman/Dunn’s).

A fraction of these suprabasal undifferentiated cells were still proliferating at E14.5 but not at E15.5 (Fig. 1C). This suggested that differentiation is not associated with upward movement at early stages.

We then asked whether cell delamination dynamics also changed as a function of developmental stage. At E14.5, detachment of cells from the basement membrane was regularly observed, with upward movement lasting around 120 minutes. This detachment was frequently associated with a cell division, either occurring after a perpendicular division or followed by a mitosis in the suprabasal layer (Fig. 1D; Supplementary Movie 1). In contrast, at E15.5 delaminations were substantially slower, lasting at least 300 minutes, and were not associated with divisions (Fig. 1E; Supplementary Movie 2). The geometry of delaminating cells also changed; at E14.5 cells rounded up for delamination, while at E15.5 cells transitioned instead into axially elongated and basally constricted, wedge-like shapes prior to delamination (Fig. 1D-F).

To better understand how these changes in division and delamination strategies relate to cell differentiation, and to identify the mechanisms involved, we turned to fixed tissue imaging. Previous work has implicated the desmosomal cadherin Desmoglein1 (Dsg1) in delamination ^15^. At E14.5 no Dsg1 or K10-positive differentiating cells were detected in the basal layer (Fig. 1G). In contrast, at E15.5 Dsg1+ K10+ cells were now observed basally, indicating that these cells had already initiated commitment towards differentiation. These double-positive basal cells exhibited enhanced localization of Dsg1 (Fig. 1H) and E-cadherin (Supplementary Fig. 1C) particularly at apical interfaces, displaying wedge-like shapes similar to delaminating cells seen in live imaging (Fig. 1H).

To precisely quantify the shape changes of these K10+-basal cells at E15.5, we performed super-resolution imaging of skin whole mounts. K10+ basal cells were still attached to the basement membrane as shown by α6β4 integrin staining, (Fig. 1I), and displayed a wedge-like shape characterized by decreased basal adhesion length and increased apical adhesion surface (Fig. 1I). Interestingly, integrins were enriched at these narrow basal footprints, indicating that differentiation is initiated prior to complete cell detachment from the basement membrane (Fig. 1J).

Collectively, these live imaging experiments are consistent with earlier work on fixed samples showing that at early developmental stages embryos engage both perpendicular cell divisions and rapid delaminations associated with suprabasal divisions to generate suprabasal layers ^6–8,10^. Our data further indicates that at E15.5 and E16.5 embryos employ a substantially slower, division-independent delamination process associated with remodeling of cell adhesion and changes in cell shape to position cells suprabasally.

### A biophysical model predicts that in the presence of a mechanical boundary, delamination depends on the mechanical properties of the delaminating cell

To understand the mechanistic basis for this switch in multilayering strategy, we turned to quantitative mechanical modelling. For this we utilized open-source 3D vertex model code ^16^ and extended a previously published 3D vertex model of the epidermis ^13^ (see Supplementary Methods). Similar models were recently used to study clonal dynamics of tumor cells in stratified epithelia ^17^ and initiation of multilayering in single-layered epithelia ^18^. The model consisted of suprabasal cells (brown), basal cells (light blue) and a basement membrane (dark blue; Fig. 2A). A key model feature was basal layer rigidity, governed by cell shape. We therefore parameterized this by Δ𝑠, the distance in shape space to the fluid-solid transition ^16,19,20^ (Fig. 2A, inset). The model also included heterotypic interfacial tensions between basal and adjacent suprabasal cells (σ_𝑎_; Fig. 2A, magenta), and a potentially wetting interfacial tension between basal cells and the basement membrane (σ_𝑏_ ; Fig. 2A green). Previous work ^13^ had shown that these three “tissue background” parameters (σ_𝑎_, σ_𝑏_, and Δ𝑠) generate a robust mechanical model that is capable of capturing key features of the multilayered tissue architecture observed *in vivo.* A central feature of this mechanical model is generation of a mechanical boundary between the basal and suprabasal layer that ensures that small active mechanical fluctuations do not induce mixing between basal and suprabasal cells, and the layers are robustly separated, as observed at E15.5 onwards.

**Figure 2.**
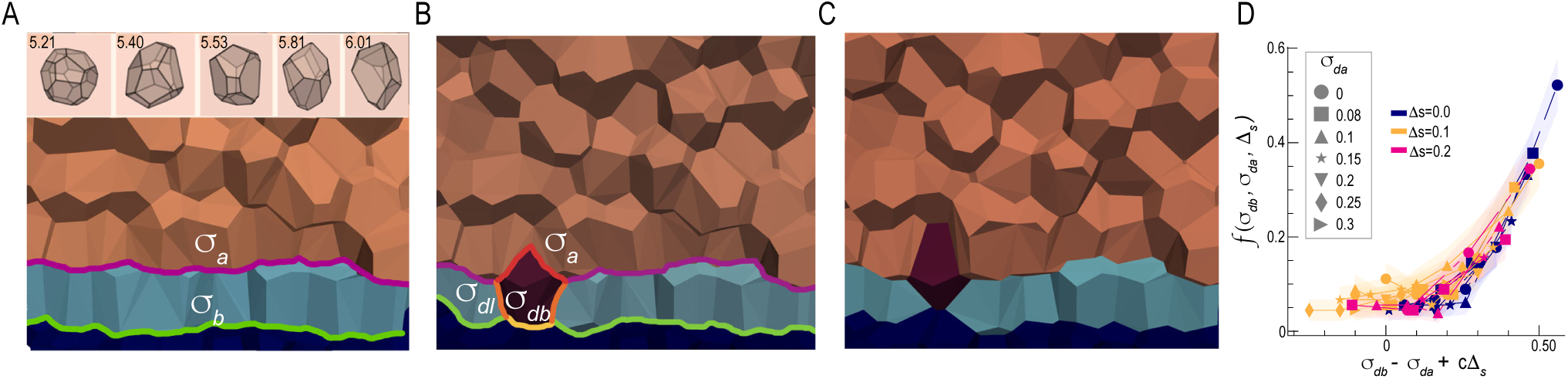
A biophysical model predicts that in the presence of a mechanical barrier, delamination depends on the mechanical properties of the delaminating cell. **A.** Snapshot from 3D vertex model for stratified epithelium geometry, illustrating suprabasal cells (brown), basal cells (light blue) and basement membrane (dark blue), interfacial tensions between the apical surface of basal cells and the adjacent suprabasal cell (, magenta), basal cells and basement membrane (σ_𝑏_, green), and cell shapes (inset) with different 3D shape parameters. **B.** Simulation snapshot including a delaminating cell type, with potentially different delaminating cell-basal cell lateral interfaces σ_𝑑𝑙_, delaminating cell-basement membrane interfaces (σ_𝑏_, yellow), and delaminating cell-suprabasal cell interfaces (σ_𝑑𝑎_, red). **C.** Snapshot of simulation showing cell in process of delaminating (cell integrates fully into the suprabasal layer by end of simulation). **D.** Fraction of cells that delaminate after a time t = X, as a function of σ_𝑑𝑏_, for many different values of the other parameters. Data averaged over an ensemble of n=20 simulations for each simulation condition. **E.** Delamination fraction collapses as a specific function of vertex model parameters.

We first sought to predict whether the presence of a mechanical boundary could explain the embryonic stage-specific differences in delamination speed and shape of the delaminating cell (see Fig, 1), similar to what has been theoretically predicted for cell extrusion in simple epithelia ^21^. To this end we specifically altered interfacial tensions of the delaminating cell – lateral σ_𝑑𝑙_, basal σ_𝑑𝑏_, and apical σ_𝑑𝑎_ (Fig. 2B). Past experimental observations ^9,22^ suggested that basal layer rigidity Δ𝑠 is also relevant for delamination. We therefore also allowed it to vary while keeping the other “tissue background” parameters σ_𝑏_ and σ_𝑎_ fixed. We found that for a cell to move upward into the suprabasal layer in the presence of small active fluctuations, it indeed needed only to change its interfacial tensions (Fig. 2C). We proceeded to precisely predict how changes in delaminating cell mechanics and the mechanical boundary itself alter delamination rates. To quantify these rates, we simulated a large ensemble of conditions (N>85) for each set of model parameters (3240 sets), and measured the fraction of cells that had delaminated in a fixed time window (Δ𝑡 = 2000 simulation time units, estimated to be 20 – 200 min ^23,24^, see Supplemental Methods). We found that delamination rates are a complex function of the four variables (σ_𝑑𝑎_, σ_𝑑𝑙_, σ_𝑑𝑏_, Δ𝑠; Supplementary Fig. 2A). To parse the resulting large parameter space, we searched for a data collapse in delamination rates ^25^, and further investigated whether the rates were consistent with an Arrhenius process governed by an energy barrier, i.e. where the rate is proportional to *e^−E/KTeff^*, where *KT_eff_* sets the magnitude of active fluctuations ^26–28^. We identified a specific combination of model parameters where all delamination rate curves collapsed on top of each other (Fig. 2D; non-collapsed data in Supplementary Fig. 2), indicating that a delaminating cell experiences a mechanical energy barrier to move upwards. Further, this barrier can be modulated by the mechanical properties of the delaminating cell, in the following form:

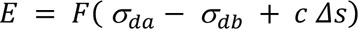

Here 𝐹 is a monotonically increasing function of its argument (see Supplemental Methods). The first two terms indicate that the energy barrier depends on the difference between the basal and apical interface tensions of the delaminating cell. To decrease the energy barrier and delaminate more efficiently, a delaminating cell needs to increase the tension (i.e. decrease the wetting) with the basement membrane, while also decreasing its apical interfacial tension with the suprabasal cells.

This is consistent with the experimentally observed emergence of a wedge-like cell shape and apical Dsg1 enrichment from E15.5 onwards (see Fig. 1G-I). The third term indicated that the energy barrier increases with increasing surrounding basal layer rigidity 𝛥𝑠. Interestingly, altering the interfacial tension σ_𝑑𝑙_ of the delaminating cell did not change the delamination rate (Fig. 2D).

Overall, these results indicated that the non-mixing of basal and suprabasal cells observed experimentally at E15.5 and E16.5 (see Fig. 1) could be explained by a mechanical boundary between these cell layers, and that cells would be able to cross this barrier by changing their own mechanical properties.

### A mechanical boundary emerges at E15.5 impacting delamination

Next, we investigated whether such a mechanical intratissue boundary emerges during development. To this end we performed detailed quantitative morphometry of basal cells across developmental stages. 3D segmentation revealed substantial transformations of cell shapes: whereas basal cells at E14.5 showed considerable shape heterogeneity and a high degree of shape deformation, at E15.5 cells shortened and became more cuboidal, and finally transitioned into homogenous, prolate cell shapes at E16.5 (Fig. 3A-C). To infer changes in interfacial tensions from these morphological transitions, we built on previous work suggesting that an apical curvature angle 𝛾 and cell height ℎ reflect a force balance along each interface ^29,30^. Further, interfacial properties – orientation angle of the lateral interface (𝛽) and the roughness of the apical interface (𝑅) - depend on heterotypic interfacial tensions ^31,32^. We therefore measured these observables *in vivo* in the basal layer (Fig. 3D- H). These measurements revealed a decrease in cell height between E15.5 and E16.5, and non-monotonic behaviors for the lateral interface orientation angle from E14.5 to E16.5 (Fig. 3E), consistent with the observed stage-specific cell shape transitions (Fig 3A-C). As in previous work, 𝑅 quantifies the tissue-scale roughness – variations between the heights of the lateral junctions across the basal layer (Fig. 3F) – which are distinct from the cell-scale curvatures defined by 𝛾 (Fig. 3G).

**Figure 3.**
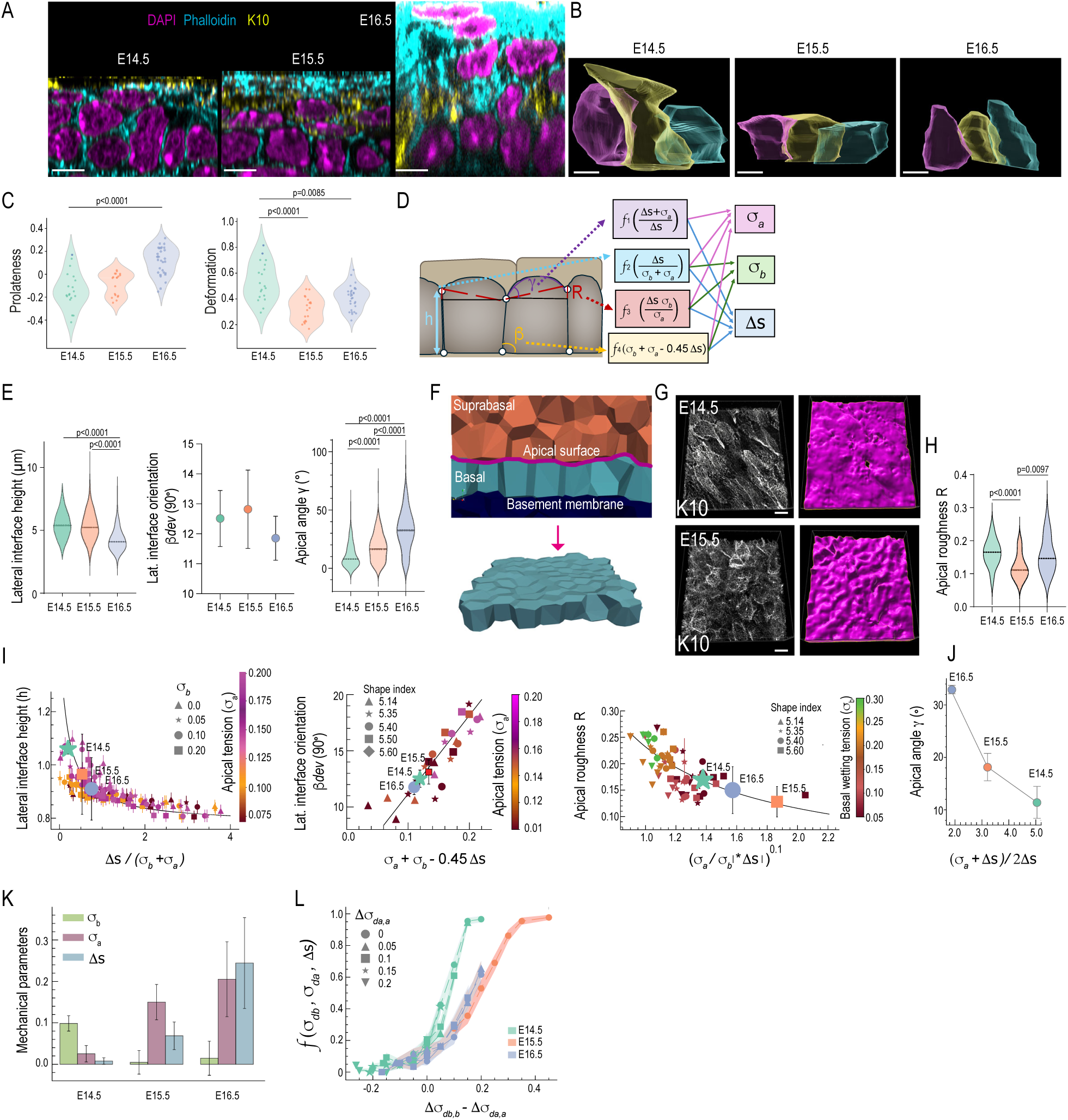
A mechanical boundary emerges at E15.5 impacting delamination. **A.** Representative super resolution images of tissue cross sections across developmental stages. Scale bars 10 µm. **B.** 3D rendering of cell shapes across developmental stages from data in (A). Scale bars 5µm. **C.** Quantification of 3D cell shapes from (B) using spherical harmonics reveals a decrease in deformed shapes and an increase in prolateness from E14.5 to E16.5 (n=19 (E14.5, E15.5)/ 30 (E16.5) cells pooled across 3 embryos; Kruskal-Wallis/Dunn’s). **D.** Schematic representation of cell- scale geometric parameters and their integration into the vertex model. **E.** Quantification of lateral interface height, variability of orientation angle and apical surface angle across developmental stages (n>400 cells (height); >25 fields of view (apical angle) >740 cells (lateral angle) per timepoint pooled across 3 mice (E14.5, E15.4), 5 mice (E15.5); Kruskal-Wallis/Dunn’s). **F.** Schematic representation of tissue-scale geometric parameter of surface roughness in the 3D vertex model. **G, H.** Representative experimental whole mount (B) and cross section images (C) of surface roughness measurements (scale bars 10 µm). **I.** Collapsed simulation data measured in the 3D model simulations across mechanical parameter space (𝜎_𝑎_, 𝜎_𝑏_, Δ𝑠). Corresponding averages of *in vivo* observables converted to simulation units specified as green, orange and blue symbols (n>85 simulations for each parameter set). **J.** Apical angle simulation data as a function of vertex model parameters required by force balance (n>85 simulations for each parameter set). **K.** Best-fit vertex model parameters 𝜎_𝑏_ (green), 𝜎_𝑎_ (pink), Δ𝑠 (blue) to the cell and tissue geometry observed at each stage; error bars represent uncertainty in the fit (see Supplemental Methods). **L.** Simulation-predicted delamination rates when delaminating cells alter their mechanical properties from the background basal cells by Δ𝜎_*da,a*_ (apical) and Δ𝜎_*db,b*_ (basal). Rate is the fraction that delaminates in a fixed time window t = 2000 timesteps, corresponding to τ ∼ 20 simulation time units (n>85 simulations per parameter set).

This can be seen by linking the images in Fig 3G to the data in Fig 3E and H - at E15.5 the cell-scale curvature 𝛾 is high but over larger length scales the apical surface is relatively flat (low 𝑅), while at E14.5 the cell-scale curvature 𝛾 is low but with a higher tissue-scale roughness 𝑅. At E16.5, 𝛾 is even higher than at E15.5, while 𝑅 is lower that E15.5. Collectively these shape changes were indicative of dynamic, stage-specific changes in apical heterotypic interfacial tension, wetting at the basement membrane, and overall tissue stiffness.

We then proceeded to integrate the experimental quantifications with the 3D vertex model by converting experimental measurements to simulation units, followed by model simulations across a wide range of parameter conditions (Fig. 3D, I, J). For three observables (𝑅, 𝛽, ℎ), we discovered that the data could be collapsed as a simple function of the model parameters (Fig. 3I; Supplementary Fig. 3.A-C). As a result, the average value of each *in vivo* observable (Fig. 3E, H) specified a point on the y-axis of the respective simulation plot (Fig. 3I), and therefore generated a unique constraint on the model parameters. Although the curved interfaces described by 𝛾 are not explicitly embedded in 3D vertex models, we can use the assumption of force balance^29,30^ to write 𝛾 directly as function of σ_𝑏_ and 𝛥𝑠 (Fig. 3J), and thereby directly extract a fourth constraint on vertex model parameters.

Collectively this benchmarking process of *in vivo* measurements to simulations resulted in a set of four constraint equations for the three unknowns (σ_𝑏_, σ_𝑎_, 𝛥𝑠) at each developmental stage (Fig. 3D); we solved this system of equations for the best-fit model parameters at each stage using a standard over-constrained solver^33^ (Fig. 3K; see Supplementary Methods).

These data demonstrated that at E14.5, the basal layer is nearly fluid-like, with very low tissue stiffness parameterized by 𝛥𝑠 (Fig. 3N). At this stage the interfacial tension between basal and suprabasal cells is low, coupled to a positive interfacial tension at the cell-basement membrane interface, suggesting only low levels of integrin-based wetting. At E15.5, the tension at the basement membrane interface became nearly zero, suggesting wetting behavior (Fig. 3K). This was accompanied by increased interfacial tension at the basal-suprabasal interface, as well as increased basal layer stiffness (Fig. 3K). At E16.5, similar trends remained, while tissue stiffness increased further (Fig. 3K). Combined, these changes generated a four-fold increase in the magnitude of the mechanical energy barrier for a basal cell to delaminate from E14.5 to E15.5 (Supplementary Fig. 3F-H).

Building on our understanding from the data collapse in Fig. 2, we were then able to collapse the rates of delamination as a function of the differences in the interfacial tensions between the delaminating cell and the background basal cells at the apical (Δ𝜎_*da,a*_) and basal (Δ𝜎_*db,b*_) interfaces (Fig. 3L; Supplementary Methods Fig. 3C, see Supplemental Methods). This analysis showed that at E14.5, the probability of a standard basal cell (0.0 on the x-axis of Fig 3L) delaminating during the time window Δ𝑡 was less than 20%, but only a small change in basal and apical tensions (∼15%, 0.15 on the x-axis in Fig 3L) was sufficient to generate a massive increase in delamination probability, increasing it to nearly 100% in the same time window. In contrast, at E15.5 and E16.5, a basal cell must change its mechanical properties substantially (by appr. 40%, 0.4 on the x-axis of Fig. 3L) in order to generate a similar increase in delamination probability.

Collectively, these data suggest that a mechanical boundary forms between the basal and suprabasal layers at around E15.5, thus requiring cells to have a robust mechanism to commit to delamination and overcome this barrier, while such a mechanism is not necessary at E14.5.

### A committed basal population characterized by Notch activation emerges at E15.5

To identify the molecular driver for this predicted mechanical state change required for delamination at E15.5 and later, we investigated the evolution of epidermal cell states using single cell RNA sequencing (scRNAseq; Fig. 4A). We obtained single-cell suspensions of skin from E13.5, E14.5, E15.5. and E16.5 embryos, isolated and sequenced in parallel to avoid batch effects (Fig. 4A). The quality-controlled and filtered dataset, obtained from 2 biological replicates/stage, comprised of >15,000 cells/stage that were annotated based on known marker genes (Supplementary Fig. 4A, B). Focusing on the epidermal (non-hair follicle) clusters, we identified cycling and non-cycling basal cells as well as suprabasal cell states (Supplementary Fig. 4C, D). Leiden clustering revealed that cell states were clustered separately according to their developmental stage, indicating that key cell states (basal and suprabasal cells) underwent substantial transcriptional changes in transcriptomes while progressing through development (Fig. 4B).

**Figure 4.**
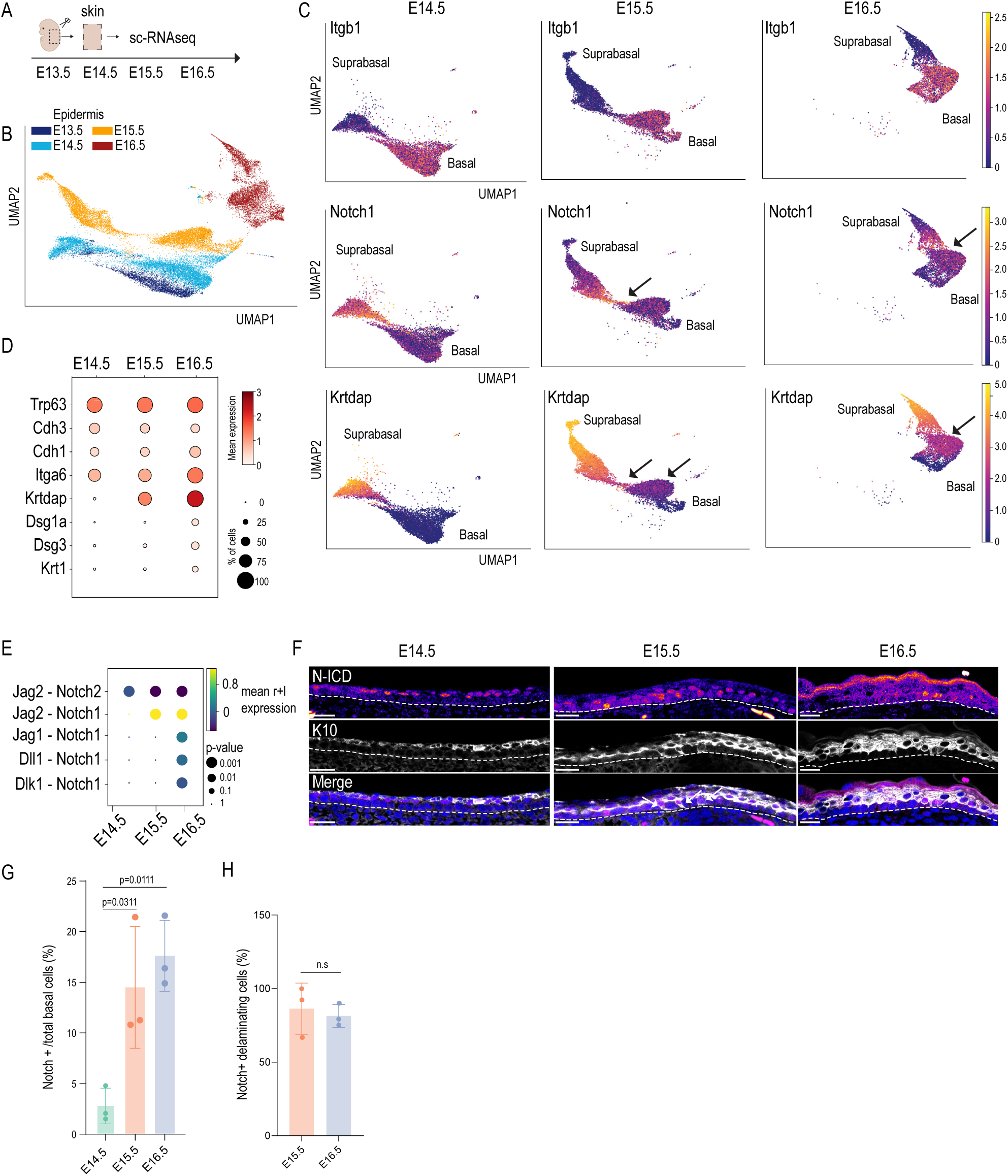
A committed basal population characterized by Notch activation emerges at E15.5. **A.** Schematics of the scRNAseq experiment. **B.** UMAP representation of the Leiden clusters of filtered epidermal cells from scRNAseq. Embryonic stages of cells are color coded. **C.** UMAP projections of basal identity and differentiation gene expression across E14.5-E16.5. Arrowheads mark transition population between basal and suprabasal cells. **D.** Dotplot of selected genes across E14.5-E16.5 basal cells. Note increased expression of differentiation genes in basal cells of E15.5 and E16.5 embryos. **E.** Dot plot of receptor-ligand (r+l) analyses between basal stem cells across E14.5-E16.5. Note increased predicted Jag2-Notch signaling interaction at E15.5 and E16.5. **F-H.** Representative images (F) and quantification (G, H) of tissue sections from E14.5- 16.5 embryos stained against Notch intracellular domain (N-ICD) and K10. Note emergence of N-ICD-K10- positive delaminating basal cells at 15.5. Dotted line marks basement membrane (mean ±SD; scale bars 20 µm; n=3 embryos/stage; Two-way ANOVA/Tukey’s (G); Mann-Whitney (H)).

Closer inspection of key differentiation genes such as *Krt1*, *Krtdap* and *Notch1* revealed that at E14.5 their expression was restricted to the suprabasal cell populations. In contrast, at E15.5 and E16.5 a subset of basal cells also showed expression of differentiation genes (Fig. 4C), in agreement with some basal cells showing K10 and Dsg1 expression at this stage (Fig.1G, H) . To conclusively identify a committed basal cell population, we examined a Harmony integrated dataset of E14.5- E16.5 epidermal cells. Leiden clustering of this integrated dataset revealed a distinct population of cells characterized by expression of both basal stem cell (e.g. *Trp63* and *Itga6*) and terminal differentiation genes (Supplementary Fig. 4E, F), indicating that only from E15.5 onward, a subset of basal cells commit to differentiation. Next to this hybrid transcriptional signature of differentiation and stemness, this subset of committed basal cells also showed enhanced expression of the suprabasal adhesion proteins E-cadherin and desmoglein (Fig. 4D), consistent with a downregulation in heterotypic interfacial tension between the committed cell and suprabasal cells predicted by the model (Fig. 2D and 3O).

As Notch activity has been shown to be critical for terminal differentiation in keratinocytes ^34^, we examined Notch pathway activity specifically in the basal layer. Receptor-ligand enrichment analyses (CellPhoneDB; ^35^ predicted increased basal activity of especially Notch1-Jagged2 signaling from E15.5 onward (Fig. 4E). Indeed, immunofluorescence analysis revealed that Notch activity (cleaved intracellular domain; N-ICD), was detected in single cells only in the basal layer of E15.5 and E16.5 epidermis, but not at E14.5 (Fig. 4F, G). Intriguingly, these Notch1-active cells were also positive for the differentiation marker K10 and displayed a wedge-shaped morphology (Fig. 4F), indicating that these cells are committed to differentiation and undergoing delamination as observed by the K10/Dsg-1 positive cells (Fig.1G, H)

Collectively these data indicated the emergence of a committed basal cell state characterized by a hybrid profile of stemness and differentiation, a change in cell shape, a switch to a suprabasal cell adhesion profile and Notch activity.

### Generation of the mechanical boundary at E15.5 is concomitant with stiffening of the basement membrane and rigidification of the basal layer

We hypothesized that maturation of the tissue and specifically the basement membrane could drive the emergence of the mechanical barrier at E15.5. Combined analysis of the scRNAseq data (Fig. 5A) and quantitative multiplex tissue imaging using Imaging Mass Cytometry ^36^, revealed an increase in basement membrane maturation and cell-matrix adhesion (Fig.5A-C; Supplementary Fig. 5A). Importantly, collagen IV, a key component of the basement membrane that is essential for its rigidification and stabilization, was increased from E14.5 to E16.5 (Fig. 5B-C), suggesting maturation of the BM from E14.5-16.5. In agreement, the hemidesmosomal α6β4 integrin was also increased during this transition, and became more and more confined to the epidermal-dermal junction (Fig. 5B, C; Supplementary Fig. 5A). Consistently, atomic force microscopy-mediated force spectroscopy confirmed a significant stiffening of the basement membrane from E14.5 to E16.5 (Fig. 5D).

**Figure 5.**
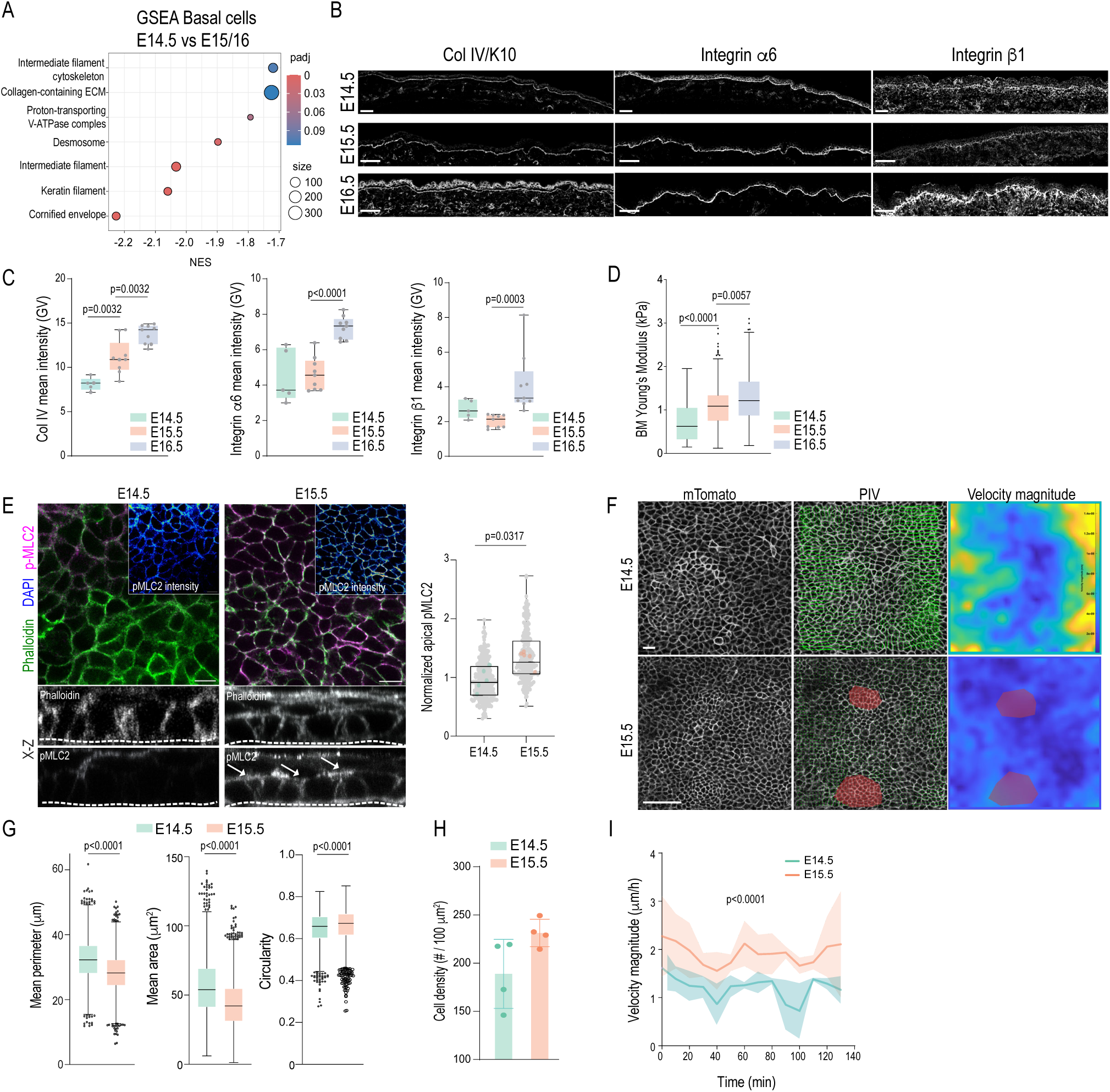
Generation of the mechanical boundary at E15.5 is concomitant with stiffening of the basement membrane and rigidification of the basal layer. **A.** Dot plot of Gene set enrichment analyses (GSEA) from differentially expressed genes in basal cells of E14.5 versus E15.5/E16.5 embryos from scRNAseq data. **B, C.** Representative images (B) and quantification (C) of tissue sections from E14.5- 16.5 embryos analyzed by mass cytometry. Note increased expression of basement membrane Collagen IV (counterstained with K10) and integrins α6 and β1 (mean ±SD; scale bars 50 µm; n=3 embryos/stage; Two-way ANOVA/Tukey’s (Left and middle panel), Kruskal-Wallis/Dunn’s (right panel)). **D.** Basement membrane Young’s moduli extracted from AFM force indentation experiments (Tukey’s box and whiskers; n>200 force curves pooled across 3embryos/stage; Kolmogorov-Smirnov). **E.** Representative images and quantification of skin wholemounts from E14.5 and E15.5 embryos stained for pMLC2 and phalloidin. Note increased pMLC2 intensity at cell junctions and interface between basal and suprabasal layers (arrows) (min-max box and whiskers; scale bars 10 µm; n=5 embryos/stage; Mann- Whitney)**. F.** Representative snapshots from live imaging movies and subsequent PIV analyses of velocity magnitudes of E14.5 and E15.5 epidermis from membrane-Tomato (mTomato) reporter mice. Red masks mark excluded hair follicle areas (scale bars 50 µm). **G.** Quantification of 2D cell perimeter, area and circularity from data in (A) (Tukey’s box and whiskers; n>2300 cells pooled across 3 embryos/stage; Mann-Whitney). **H.** Quantification of cell densities from data in (A) (mean ±SD; n=4 embryos/stage; Mann-Whitney). **I.** Quantification of velocity magnitudes from data in (A). Note attenuated velocity at E15.5 (mean ±SD; n=4 embryos/stage; Linear Mixed-Effects model.

Based on observations in other systems that link changes in tension to mechanical boundary formation ^9,16,19,20,22,37,38^ we speculated that this change in tissue mechanics promoted establishment of a mechanical boundary between basal and suprabasal cells. Quantification myosin phosphorylation (pMLC2) indicated increased actomyosin tension at apical cell-cell junctions and specifically at the interface between basal and suprabasal cells at E15.5 compared to E14.5 (Fig. 5E), providing further support for the geometry-based tension inference indicating emergence of a mechanical boundary (Fig. 3N). Furthermore, analyses of cell shapes and tissue dynamics from live imaging showed that the cells in the basal layer of E14.5 embryos were larger and more irregular shaped compared to cells at E15.5, as already seen in fixed tissue imaging (Fig. 5F, G; Fig. 3 A-C; Supplementary Movie 3). In agreement, at E15.5 cell density was increased, (Fig. 5F, H), with decreased mobility of the basal layer cells as quantified by particle image velocimetry (Fig. 5F, I; Supplementary Movie 3), in line with previous *in vitro* work implicating a jamming transition in initiation of delamination ^9^. Together, these changes were consistent with a tissue-scale rigidity transition in the developing epidermis that occurs from E14.5-16.5, from an initially floppier basement membrane and a fluid-like basal layer to a stiff basement membrane and a jammed basal layer with enhanced cell-matrix adhesion and apical actomyosin tension.

### Tissue rigidity and cell shape changes drive Notch-dependent delamination and control of tissue crowding

We next investigated whether the changes to tissue rigidity are sufficient to induce basal layer jamming and the emergence of a mechanical barrier. For this we mimicked tissue maturation *in vitro* by modulating cell density and substrate rigidity by engineering micropatterned adhesive islands on polyacrylamide hydrogels of defined stiffness (Fig. 6A). Culturing primary epidermal progenitors on micropatterned islands revealed that increasing substrate stiffness under confinement not only substantially enhanced differentiation (Fig. 6B), but, importantly, promoted efficient suprabasal separation of these differentiated cell populations from the undifferentiated cells that remained basally positioned (Fig. 6C), indicating that increasing substrate stiffness is sufficient to induce boundary formation.

**Figure 6.**
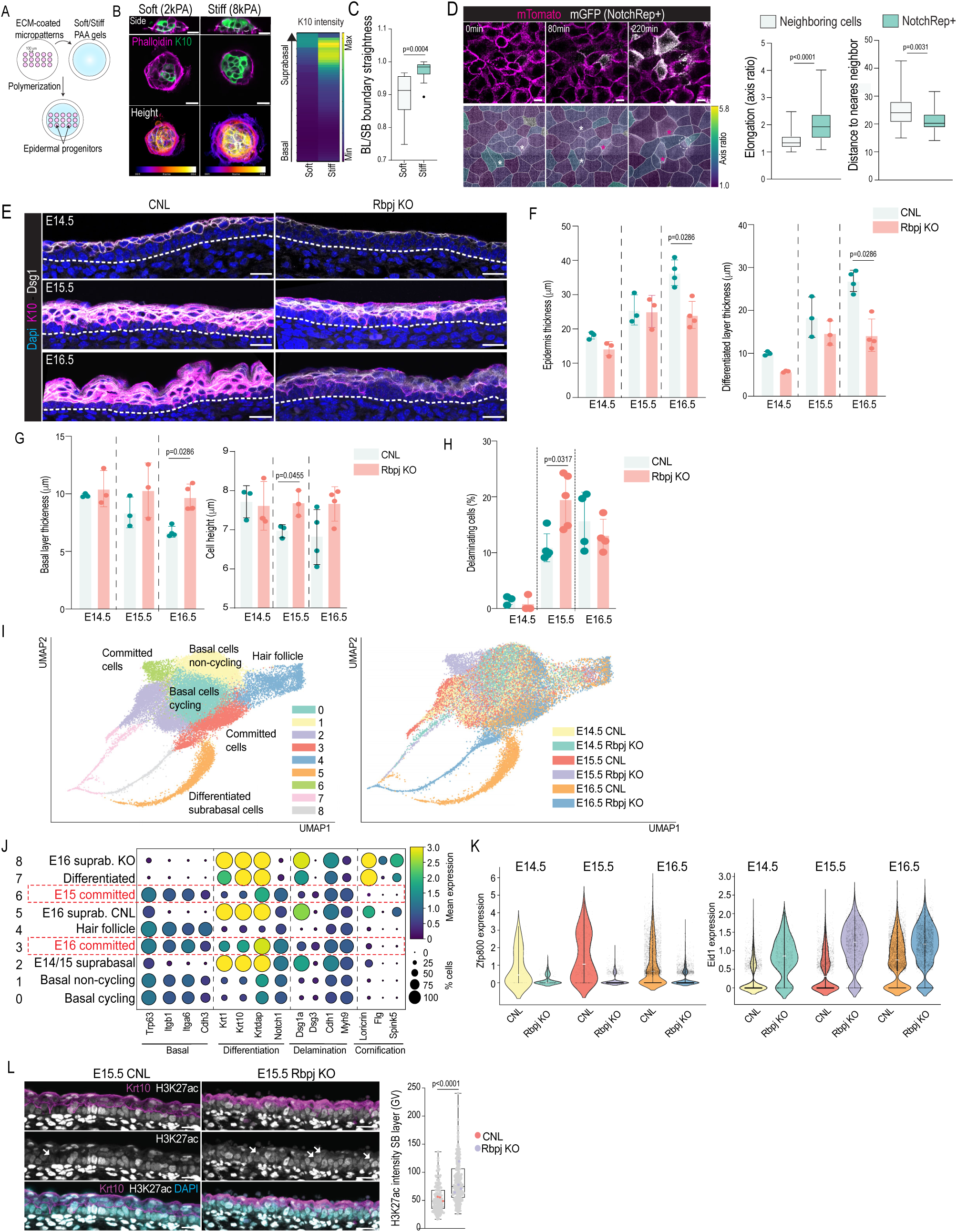
Tissue rigidity and cell shape changes drive Notch- delamination and control of tissue crowding. **A.** Schematic illustration of micropatterning of soft and stiff polyacrylamide (PAA) hydrogels. **B.** Representative images of primary epidermal progenitors stained with K10 (green) and phalloidin (magenta) cultured on 100-µm-diameter micropatterns on 2.5 kPa (soft) or 8 kPa (stiff) PAA gels. K10 mean intensity was quantified as a function of the distance from the bottom of the micropatterns. Note the increased K10 mean intensity in the suprabasal region on stiff substrates. **C.** Quantification of boundary straightness between the basal layer (bottom) and the first suprabasal layer (upper layer). (Tukey’s box and whiskers, n= 10 (soft); 15 (stiff) patterns pooled across 3 independent experiments; Mann-Whitney). **D.** Representative snapshots and quantification of live imaging movies from epidermal progenitor cell monolayers from mTomato/mGFP-Cre reporter mice crossed with Notch1- ICD-Cre to detect cells that activate Notch (mGFP+). Note increased elongation and local cell density of cells (white asterisk) that subsequently activate Notch signaling (red asterisk). Dotted circle marks delamination of Notch-Reporter-positive (NotchRep+) cell (min-max box and whiskers; n=38 Notch+ cells pooled across 3 independent experiments; Mann-Whitney; scale bars 10 µm). **E.** Representative images of tissue sections from E14.5-16.5 embryos of control and Rbpj-KO mice stained against K10 (magenta) and Desmoglein 1 (Dsg1; white) (images representative of >3 embryos/stage; scale bars 20 µm). **F.** Quantification of epidermal thickness from data in (A). Note reduced total and differentiated layer thickness of Rbpj-KO epidermis at E16.5 (mean ±SD; n=3 embryos/E14.5 and E15.5, 4/E16.5; Mann-Whitney). **G.** Quantification of basal layer thickness and cell height from data in (E). Note increase basal layer thickness and cell height of Rbpj-KO epidermis emerging at E15.5 (mean ±SD; n=3 embryos/E14.5 and E15.5, 4/E16.5; Mann-Whitney). **H.** Quantification of delaminating cells from data in (E). Note increase in delaminating cells in Rbpj-KO epidermis at E15.5 (mean ±SD; n=3 (E14.5), 5 (E15.5), 4 (E16.5) embryos; Mann-Whitney). **I.** UMAP representation of the Leiden clusters (left panel) and biological samples (right panel) of filtered epidermal cells from scRNAseq. **J.** Dotplot of selected genes across E14.5-E16.5 epidermal cells. Clusters 3 and 6 (in red) represent committed basal cell populations enriched in Rbpj-KO cells of E15.5 and E16.5 embryos. **K.** Violin plots of Zfp800 of Eid1 expression across E14.5-E16.5 in control and Rbpj-KO epidermal cells. **L.** Representative images and H3K27ac nuclear intensity quantification from tissue sections from E15.5 embryos of control and Rbpj-KO mice stained for H3K27ac (grey), K10 (magenta) and DAPI (cyan) (n= 176 (CNL), 234 (Rbpj-KO) cells pooled across 3 (CNL) /4 (Rbpj-KO) embryos; Student’s t-test; Scale bars 20 µm).

As the emergence of basal committed cells was associated with Notch activity (Fig. 4D), we then asked whether basal layer crowding was sufficient to induce Notch signaling. For this, we performed live imaging of primary epidermal progenitors isolated from Notch1 activity reporter mice ^39^ to identify whether and when Notch signaling is activated in basal cells triggered to differentiate and form a suprabasal layer (Fig. 6D). As expected, all progenitors were initially Notch-reporter negative, but upon application of calcium to trigger differentiation and multilayering, individual cells gradually switched on the reporter followed by delamination (Fig. 6D; Supplementary Movie 4). Intriguingly, prior to turning on Notch1 signaling these cells were more elongated and resided in denser tissue regions than reporter negative-cells (Fig. 6D), thus indicating that crowding and increased cell-cell to cell-matrix contact area promoted Notch activation.

We then asked whether Notch1 transcriptional activity is necessary for cell fate commitment and delamination by deleting the Notch transcription factor, Rbpj, in the epidermis (Rbpj-KO; ^40,41^). At E14.5, the epidermis of Rbpj-KO mice appeared largely comparable to control littermates (Fig. 6E). At E15.5, and more prominently at E16.5, the epidermis became structurally more abnormal, characterized by a specific thinning of the suprabasal differentiated layers (Fig. 6F). Moreover, basal cells were increased in height and became more columnar in E15.5 and E16.5 Rbpj-KO mice compared to controls (Fig. 6G), accompanied by higher cell density (Fig. 6D), indicating increased crowding of the basal layer. To examine if this increased crowding was due to reduced commitment to differentiation or decreased delamination, we quantified differentiating, delaminating cells. We observed emergence of the committed basal cell population in both control and Rbpj-KO epidermis (Fig. 6H), indicating that the initial commitment to differentiate does not depend on Notch transcriptional activity. Instead, we observed an increase in the number of wedge-shaped committed cells undergoing delamination at this stage (Fig. 6H), suggesting an inability to overcome the mechanical energy barrier and move suprabasally.

To identify the molecular mechanism by which Notch transcriptional activity regulates efficient delamination, we performed scRNAseq analysis across E14.5-E16.5 from control and Rbpj-KO mouse skin. Overall, basal cell populations were transcriptionally similar between controls and Rbpj- KO mice (Fig. 6I). However, as described previously ^34^, Rbpj-KO cells of all developmental stages displayed impaired terminal differentiation (Fig. 6I; Supplementary Fig. 6A). Intriguingly, at E15.5 a distinct Leiden cluster representing committed basal cell population emerged, mainly composed of Rbpj-KO cells (Fig. 6I, J; Supplementary Fig. 6B). Consistently, at E16.5 the Leiden cluster consisting of committed cells was dominated by Rbpj-KO cells (Fig. 6I, J; Supplementary Fig. 6B), validating the immunofluorescence observations that Rbpj-KO still commit to initial differentiation but cannot efficiently delaminate, leading to accumulation of committed basal cells.

To identify master transcriptional switches downstream of Notch that drive the final step of delamination, we performed differential gene expression analyses of committed cells between control and Rbpj-KO at E15.5 and E16.5. This analysis identified two interesting significantly altered regulators in Rbpj-KO committed cells: the zinc finger protein Zpf800, a master regulator of intestinal stem cell differentiation ^42^, was not upregulated (Fig. 6K left panel; Supplementary Fig. 6C, D). At the same time Rbpj-KO basal cells failed to downregulate the Ep300 Inhibitor of Differentiation (Eid1) (Fig. 6K, right panel). Eid1 acts as a transcriptional repressor through inhibition of p300-mediated H3K27 acetylation (H3K27ac), thus preventing the activation of enhancers that promote differentiation ^43,44^. Indeed, RNA *in situ* hybridization and quantitative immunofluorescence confirmed that while control embryos downregulated Eid1 in the suprabasal layers and, as a consequence, showed substantially reduced H3K27ac levels in delaminating cells and suprabasally, Rbpj-KO embryos maintained both high Eid1 and H3K27ac in delaminating cells and suprabasal layers, in line with reduced differentiation (Fig. 6L).

Collectively, these data demonstrate that stiffening of the substrate is sufficient to drive basal layer jamming and the formation of a mechanical barrier between basal and suprabasal compartments. This basal layer jamming and crowding then triggers Notch signaling required for completion of delamination and control of basal layer density.

## Discussion

Here we show that the developing epidermis employs different strategies for multilayering that are driven by changes in tissue mechanics. Early during development of this stratified epithelium, there is no substantial mechanical energy barrier between basal and suprabasal cell layers. This allows fast upward movement of basal cells independent of basal fate commitment by either dividing perpendicularly or detaching quickly in order to rapidly establish a stratified tissue. Similar mechanical-instability-driven mechanisms have been proposed by previous modeling ^18,21^. At later stages (E15.5 and beyond), maturation and stiffening of the basement membrane enhance basal cell-matrix adhesion and crowding to rigidify the tissue, generating a substantial mechanical boundary between basal and suprabasal cells. In order to cross this boundary, cells from this stage on must now commit to differentiation and undergo subsequent transcriptional rewiring, controlled by Notch activation, to complete delamination. Interestingly, this delamination process in the matured epidermis is substantially slower than the rapid detachment observed by us and others in early embryonic development^9,10^. As Notch activation itself is regulated by tissue rigidification and crowding, as also found in other systems^45,46^, it suggests an elegant control mechanism for homeostasis in an open niche not characterised by clear geometrical cues: tissue crowding itself is sufficient in a rigidified tissue to trigger Notch activation necessary to control the rate and timing of delamination. This feedback loop thus allows accurate control over the densities of both basal and suprabasal layers in coordination with compartmentalization of stem cells and differentiated progeny.

Many open questions remain. A key open question is how the system decides which subset of cells in the basal layer becomes committed and Notch active. Previous work has suggested that since Notch is a surface receptor, its activity depends on tissue crowding together with associated cell shape and surface area changes that regulate the density of such receptors ^46–48^. This could explain the emergence of Notch active cells in elongated cells within dense regions – these cells would have transiently maximized surface area to receive Notch ligand signal from their neighbors. Previous work has investigated a 2D vertex model where cell fate decisions are coupled to cell areas ^49^, which may be a useful starting point for predicting features we observe here. Future modeling work should focus on understanding how changes to tissue stiffness and cell shape induce Notch signaling in a ‘correct’ subset of basal cells (e.g. they are spaced out and the right fraction depending on the degree of crowding), and on adding the effects of differently oriented cell divisions in basal layer — in different developmental stages – to the 3D model.

Downstream of Notch signaling, it will be intriguing to understand how the transcriptional reprogramming, involving Eid1and Zfp800 associated with global changes in H3K27ac translates to control of delamination timing. It is interesting to note that the direct downstream targets of Eid1- the histone acetyltransferases P300 and CBP - are evolutionarily conserved maintenance factors of lineage-specifying core-transcriptional networks. They function as transcriptional rheostats that control global H3K27ac levels, where the duration of their activity determines the long-term impact on chromatin state ^50,51^. Upon their inhibition, >8h of hypoacetylation is required to trigger a permanent switch and “point of no return” to H3K27me3-mediated chromatin silencing ^50^.

Intriguingly, this is the approximate duration of the slower delamination process after E15.5. Thus, Notch may function as a mechanical sensor to ensure that terminal differentiation is not initiated prematurely in areas of low basal cell density. Precise coordination of onset of terminal differentiation with basal stem cell layer density is likely to be a critical feature in dynamically self-renewing tissues with an “open stem cell niche”. Collectively, we propose that in such tissues mechanical boundaries are required to separate biological activities of cycling stem cells and post-mitotic terminally differentiated cells. Here, precisely controlled cell state transitions control the transit between the compartments to robustly maintain tissue homeostasis and barrier integrity.

## Methods

### Mice

Membrane-targeted Tomato reporter mice (R26R^mT/mG^), nucleus-targeted Tomato reporter mice (R26R^nT/nG^) and high-activity Notch1 activation-dependent reporter mice (N1IP: CreHI) were from were from JAX laboratories (stock #007676, #:023035 and #027234 respectively). Rbpj floxed mice were obtained from JAX (stock # 034200) and crossed with the K14-Cre line ^40^ to obtain epidermis- specific depletion. Embryos were staged based on vaginal plug (taken as embryonic day 0) and limb morphology. All mouse studies were approved and carried out in accordance with the guidelines of the Ministry for Environment, Agriculture, Conservation and Consumer Protection of the State of North Rhine-Westphalia (LANUW; 81-02.04.2022.A065; and 2024-155), Germany.

### Ex vivo embryo imaging

Imaging was performed essentially as described earlier ^9,13^. Briefly, embryos were dissected at embryonic stages E14.5 or E15.5 and immobilized on custom-built imaging chambers engineered on top of Lumox-teflon imaging plates to allow gas exchange (Sarstedt). Whole embryos were immersed in epidermal growth medium (CnT-PR, CELLnTEC Advanced Cell Systems) supplemented with calcium-depleted 10% fetal bovine serum (HyClone/Thermo Fisher), 20 U/ml penicillin-streptomycin (Thermo Fisher), 20 mM Hepes, 1.8 mM CaCl2, and 10 ng/ml mouse recombinant EGF (Novus; NBP2-34952). Imaging was performed using an Andor Dragonfly 505 high-speed spinning disk confocal microscope (Oxford Instruments) with an inverted Nikon Eclipse Ti2 microscope (Nikon) equipped with 488 nm and 546 nm lasers, an Andor Zyla 4.2 sCMOS camera and an environmental chamber set at 37°C, 5% CO2 using a 40X dry objective. Images were acquired using Fusion 2.0 software (Andor). After acquisition, the image series were 3D drift corrected using the ImageJ plug-in Fast4Dreg ^52^ combined with manual registration, if required.

### Immunofluorescence and confocal microscopy

For skin whole mounts, embryos were dissected from pregnant mice and directly fixed in pre- warmed (37°C) 4% paraformaldehyde (PFA) at room temperature for 3 h. After multiple PBS washes, the skin was carefully peeled using forceps and placed in blocking solution (0.5% Triton-X, 5% bovine serum albumin (BSA), 3% normal goat serum (NGS)) for 1 h at room temperature. For paraffin sections tissue biopsies were fixed in 4% PFA, embedded in paraffin and sectioned. Sections were de-paraffinized using a graded alcohol series and antigen retrieval was carried out using target retrieval solution (Dako) at pH 6 or pH 9 in a pressure cooker. For cryosections, tissue samples were placed unfixed into cryomolds, embedded in OCT (Tissue Tek) and allowed to harden on dry ice. 8 μm thick sections were cut, air-dried and fixed with 4% PFA. For both paraffin and cryofixed sections, samples were blocked in 5% BSA, 3% NGS. For paraffin sections antibodies were diluted in Dako antibody diluent (Agilent). For cryosections antibodies were diluted in blocking solution.

Cultured epidermal cells were fixed in 4% PFA for 15 min at room temperature After PBS washes, samples were placed in 0.3% Triton-X, 5% BSA for one hour at room temperature after which primary antibodies were diluted in the same solution. In all cases antibodies were incubated overnight in moist chamber at 4°C. Following PBS washes, secondary antibodies were diluted in 1% BSA in PBS and incubated for 1h at room temperature.

For super-resolution imaging of Actin (Phalloidin), K10, Dsg1, E-cadherin, and Integrin α6, the embryos were fixed in pre-warmed 4% PFA at 37°C for 1 hour and rinsed with PBS. The back skin was peeled using forceps and incubated in blocking solution (2% Triton-x, 5% BSA in PBS) for 24 hours at 37°C. Primary antibodies were diluted in the blocking solution and incubated 48 hours at 37°C on shaker followed by several washes with 2% Triton-X with a final overnight wash.

Secondary antibodies were diluted in blocking solution and incubated 24 hours at 4°C followed by several washes with 2% Triton-X.

For pMLC2 staining embryos were fixed in 4% PFA dissolved in HBSS for 1 hour on a shaker at room temperature. Back skins samples are then peeled and permeabilized with 0.02% PBS-Tween 20 for 30mins and blocked with blocking solution (5% NGS,1% BSA) for 2 hours before incubating with primary antibody overnight in 4⁰C. Samples are then thoroughly washed with 0.02% PBS- Tween 20, incubated with secondary antibodies at 1:400 dilution for 1 hour at room temperature and washed. All tissue samples were mounted in Elvanol.

The following primary antibodies were used: guinea-pig anti-Keratin 10 (Progen, #GP-10; 1/300); rabbit anti-p-MLC2 (p-Ser19; Cell Signaling, 3675; 1:200), rabbit anti-cleaved-Notch1 (Cell signaling; #4147S; 1/50), mouse anti-Desmoglein 1/2 (Progen, #61002; 1/200), rabbit anti-TGM1 (360; Proteintech, #12912-3-AP; 1/300), rabbit anti P63 (Genetex #GTX102425; 1/100), guinea-pig anti-Keratin 14 (Progen, GP-CK14; 1:300), mouse anti E-cadherin (BD Biosciences, #610181; 1/200), rabbit anti Cyclin B1 (Cell Signaling, #4138S; 1/200), mouse anti H3K27ac (Active Motif, #39685, 1/200), rat anti integrin α6 (Biorad #MCA699; 1/200) . F-actin was detected using Alexa- Fluor-conjugated phalloidin (1:500). Bound primary antibody was detected by incubation with Alexa-Fluor 488-, 568- or 647 -conjugated antibodies (Invitrogen). Nuclei were counterstained with 4′, 6-diamidino-2-phenylindole (DAPI, Invitrogen). Slides were mounted in Elvanol. Fluorescence images were collected by laser scanning confocal microscopy (LSM980; Zeiss) with with Zeiss ZEN Microscopy software (Zeiss ZEN version 3.7) using a 40X water immersion objective. Images were acquired at room temperature using sequential scanning of frames of 1 µm thick confocal planes (pinhole 1). Images were collected with the same settings for all samples within an experiment. For super-resolution imaging a 63X oil objective and the Airyscan 2 SR (super- resolution) mode with default resolution settings were used. Laser power and image size were kept at a minimum to avoid photo-bleaching. All airyscan images were 3D processed using the Zeiss Airyscan Processing mode with auto filter and standard strength.

### Embryonic epidermal vertex analysis

Membrane-tomato positive E14.5, E15.5 and E16.5 embryos were isolated, fixed immediately in 4%PFA over night at 4°C and washed in PBS. The back skin was then removed using micro-scissors and forceps and permeabilized in PBS/0.5% Triton-X for 1h at RT. Nuclei were stained by incubating the skin in DAPI diluted in DAKO antibody diluent for 1h at RT. The skin was then mounted in Mowiol.

Confocal stacks were recorded at a voxel size of X: 0.361, Y: 0361, Z:0.299 using a Leica SP8 equipped with an APO CS2 63x/1.40 oil objective. To determine basal cell vertices, confocal stacks were processed in Imaris and minimal XZ or YZ projection were exported using the extended section view. Using the segmented line tool in ImageJ a line was drawn across the apical and basal vertices of basal cells. Coordinates were extracted and “Julia” was used in Visual Code Studio to determine cell height (h), apical and basal vertex distances (a) and the lateral interface orientation (angle beta). The lines connecting the basal and apical vertices, respectively, were used to determine the apical layer roughness (R) (See also Supplementary Methods). The apical curvature angle gamma was determined manually using the angle tool in ImageJ.

### Image analyses

For tissue-scale analysis, 2D segmentation were performed from live imaging data on whole embryos or cultured epidermal progenitor monolayers using Cellpose4SAM (v4.0.8) ^53^. 2D cell shape parameters were measured using custom Python script and cell densities were measured using ImageJ. For 3D analyses of cell shapes of delaminating cells from embryo live imaging, Imaris 11 was used to manually create 3D surfaces of delaminating and corresponding neighboring basal cells. For 3D segmentation of basal cells from super resolution images, K10 negative basal cells were manually segmented in 3D using Imaris. Meshes were exported from Imaris surfaces after which shapes were analyzed by quantifying Spherical Harmonics ^54^ using a custom Python script.

In fixed tissue samples, basal committed cells and delaminations were identified based on the expression of specific markers (K10 and/or Desmoglein) and their contact with the basement membrane, and subsequently manually counted. Active-Notch positive cells were quantified manually. The method was applied across all stages and conditions. Integrin α6, E-cadherin and Desmoglein mean intensities were manually measured at the basal, apical and lateral membranes on Image J. To quantify epidermis thickness, the distance between epidermal basement membrane and most superficial point of the epidermis was measured. To quantify specifically the basal layer thickness versus the differentiated layer thickness, the expression of specific markers (K10 and desmoglein) has been used to distinguish the differentiated layers (K10+) from the basal layer.

To measure boundary straightness between the bottom and upper layers in micropatterns of soft and stiff polyacrylamide (PAA) hydrogels, both a segmented line following the top of each nucleus in the bottom layer and a straight line were drawn. The lengths of these lines were measured using ImageJ, and the ratio of the straight line to the segmented line was calculated.

Eid1 RNA signal was quantified by segmenting suprabasal cells based on Keratin-DAP–positive cells. Mean fluorescence intensity of Eid1 mRNA was measured using Fiji.

### 3D vertex model

A dynamic three-dimensional model of stratified epithelial tissues was developed using the 3D vertex framework (Zhang and Schwarz, 2022). In the 3D vertex model, cells are represented as polyhedra, each characterized by a homeostatic volume 𝑉_0_ and surface area 𝑆_0_. The geometry of each cell is defined by its vertices, which serve as the degrees of freedom of the system. The modeled epithelium consists of two main cell compartments. The upper compartment represents the suprabasal differentiated cell layer, while the middle one-cell-thick layer corresponds to the basal stem cell layer. Because the vertex model treats cells as space-filling polyhedra, we modeled the basement membrane as an additional layer of interconnected “cellular network” directly attached beneath the basal layer. In this framework, cells exert forces on one another through shape-based interactions governed by the following energy functional :

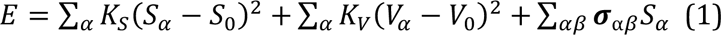

The first term of the energy functional describes the competition between contractile forces generated by cell actomyosin networks (which enable cells to minimize their shape or surface area to resist their environments), and adhesive forces mediated by cadherin-based, which promote the elongation of cell– cell contacts during neighbors’ exchange (T1s). The second term accounts for the mechanical resistance of cells to volume fluctuations, as most cells are fluid-filled and resist changes in volume. The third term incorporates heterotypic interfacial tension 𝜎_𝑎_ between the basal cells and suprabasal cells, and the tension 𝜎_𝑏_ associated with integrin-based wetting behaviors at basal-basement membrane interface.

Previous studies have shown that differential expression of adhesion molecules and cytoskeletal components associated with cell fate specification (through variations in protein or gene expression) can modify cell–cell interactions, leading to differences in adhesion strength and heterotypic interfacial tension ^9,13,55–63^. Such tension differences are known to play a central role in establishing and maintaining sharp tissue boundaries ^61^. Moreover, basal cells adhere to the basement membrane via integrins, which introduce a wetting interaction characterized by 𝜎_𝑏_^64^. The parameters 𝐾_𝑆_ and 𝐾_𝑉_ denote the elastic moduli associated with the surface area and volume of the cells, respectively. Upon non-dimensionalization, this energy functional allows the definition of a key order parameter known as the target shape index, 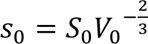, which controls the mechanical rigidity of the tissue. By tuning 𝑠_0_, the epithelial confluent tissue exhibits a rigidity transition from a solid-like to a fluid-like state, with a critical value at 𝑠_0_ = 5.4 ^16,20^.

To quantitatively predict the mechanical parameters and cellular mechanisms underlying mechanical barrier formation between the basal and suprabasal layers, we first simulated the epithelium in the absence of cell delamination. The system evolves according to overdamped Langevin dynamics, where vertex positions are continuously updated to minimize the total mechanical energy:

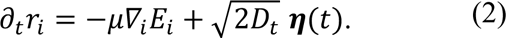

The left-hand side represents viscous drag forces, the first term on the right-hand side corresponds to interaction forces derived from the energy potential, and the second term represents thermal fluctuations in vertex positions 𝑟_𝑖_ . Here 𝜼 is an uncorrelated stochastic Gaussian white noise with unit amplitude, 𝐷_𝑡_ = 𝜇𝑘_𝐵_𝑇_788_, where 𝜇 is the mobility (inverse frictional drag), 𝑘_𝐵_ Boltzmann constant, and 𝑇_788_ is an effective temperature parameter that sets the scale of the fluctuations. Unless otherwise noted, for our simulations we choose 𝜇 = 1, 𝑘_𝐵_ = 1 and 𝑇_788_ = 0.002. This choice of 𝑇_788_ allows fluctuations of interface lengths but is not sufficient to generate diffusion or fluidity in the basal layer.

From the simulated model, we can compute the effective tensions governing different interfaces: a) The homotypic tension between basal cells 𝜎_𝑏𝑏_ = 4𝐾_𝑆_Δ𝑠_𝑏_ where Δ𝑠 = 𝑠_𝛼_ − 𝑠_0_ measures the basal layer stiffness (and Δ𝑠 = 0 at the transition point); b) The effective heterotypic tension at the basal-suprabasal interface 𝛤_𝑎_ = 2𝜎_𝑎_ + 2𝐾_𝑠_(Δ𝑠_𝑠_ + Δ𝑠_𝑏_), where Δ𝑠_𝑠_the stiffness of the suprabasal layer; c) The effective wetting tension 𝛤_𝑏𝑚_ =2 𝜎_𝑏_ +2𝐾_𝑠_(Δ𝑠_𝑏_ + Δ𝑠_𝑚_) between basal cells and the basement membrane (constituted of a cellular network with a small shape parameter). Unless otherwise noted, Δ𝑠_𝑚_ = 0.26, which ensures the basement membrane remains very stiff and solid-like.

For more detailed description see Supplementary Methods.

### Single-cell RNA sequencing (scRNAseq)

E13.5, E14.5, E15.5 and E16.5 embryos from wild type, control or K14Cre-RBP-J floxed/floxed mice (two embryos/genotype/stages) were collected simultaneously and back skins were peeled off. Single-cell suspensions were generated by enzymatic digestion by incubating back skin tissues in a mixture of DNAse I (40 µg/ml), Liberase (0.125 mg/ml; Roche Cat. NO. 05401119001), and papain (30 U/ml; Worthington) in Minimum Essential Medium, Spinner modification (Gibco 11380) for 15 min (E13.5), 20 min (E14.5), 30 min (E15.5) or 40min (E16.5) at 37°C followed by mechanical dissociation. The reaction was stopped with 22% FBS and single cells were washed with FACS buffer (0.2% chelated FBS, 2 mM EDTA in 1x DPBS (Sigma Aldrich, D8537)). For wild type sequencing, cells were labeled with BD Mouse Single-Cell Multiplexing Kit (Cat # 633793): E13.5 rep 1: Sample Tag (ST) 1; E13.5 rep 2: ST 2; E14.5 rep1: ST 3; E14.5 rep2: ST 4; E15.5 rep1: ST 5; E15.5 rep2: ST 6; E16.5 rep1: ST 7, E16.5 rep2: ST 8. For K14Cre-Rbpj floxed and control sequencing, surface biotinylation was used ^65^. Briefly, cells were labeled with 50 µg/mL EZ-Link sulfo-NHS-biotin (Thermo Fisher Scientific, 21217) on ice for 10 min, followed by incubation in 0.6 µg/ml TotalSeq-A PE streptavidin hashtag antibodies (BioLegend) for 30 min on ice as follows: E14.5 Rbpj-KO 1: TotalSeq-A0951 PE Streptavidin 405251 ; E14.5 Rbpj-KO 2: TotalSeq™-A0952 PE Streptavidin 405253; E14.5 control 1: TotalSeq-A0953 PE Streptavidin 405255; E14.5 control 2: TotalSeq-A0954 PE Streptavidin 405257; E15.5 Rbpj-KO 1: TotalSeq-A0951 PE Streptavidin 405251 ; E15.5 control 1: TotalSeq-A0953 PE Streptavidin 405255; E15.5 control 2: TotalSeq- A0954 PE Streptavidin 405257; E16.5 Rbpj-KO 1: TotalSeq™-A0951 PE Streptavidin 405251 ; E16.5 Rbpj-KO 2: TotalSeq-A0952 PE Streptavidin 405253; E16.5 control 1: TotalSeq-A0953 PE Streptavidin 405255; E16.5 control 2: TotalSeq-A0954 PE Streptavidin 405257. Labeled cells were loaded on a microwell cartridge of the BD Rhapsody Express system (BD) following the manufacturer’s instructions. Single-cell whole transcriptome analysis libraries were prepared according to the manufacturer’s instructions using BD Rhapsody WTA Reagent kit (633802; BD), except for the Rbpj-KO and control sample tag libraries in which totalseq-ADT-oligo1 (5’- TGCTCTTCCGATCTTGGCACCCGAGAATTCCA-3’) primer in place of Sample Tag PCR1 and Sample Tag PCR2 primers during the Sample Tag library amplification ^65^. Libraries were sequenced on the Illumina NextSeq 500 using High Output Kit v2.5 (150 cycles; Illumina) for 2 × 75-bp paired-end reads with 8-bp single index aiming sequencing depth of >20,000 reads per cell for each sample.

### scRNAseq analyses

#### Read Data QC and Mapping

Illumina sequencing results were demultiplexed and converted to FASTQ format using Illumina’s bcl2fastq software (version 2.20.0.422). For K14Cre-RBP-J floxed/floxed data a custom demultiplexing script was used to assign barcodes to sample tags (rhapsody-demultiplex with parameters ‘-w BD_CLS1_v2.txt BD_CLS2_v2.txt BD_CLS3_v2.txt -a TGGCACCCGAGAATTCCA <fastq files> sample_tags.txt’, and sample tags as described above).

Preprocessed read data was aligned to the mouse reference genome (GRCm39) and counted with STAR ^66^ (version 2.7.11b) through its single-cell functionality STARsolo, using the following parameters: WT: ‘--runThreadN 16 --soloType CB_UMI_Complex --soloCellFilter None --outSAMtype BAM SortedByCoordinate --soloFeatures Gene GeneFull GeneFull_Ex50pAS --soloAdapterSequence NNNNNNNNNGTGANNNNNNNNNGACA --soloCBmatchWLtype 1MM --soloCBposition 2_0_2_8 2_13_2_21 3_1_3_9 --soloUMIposition 3_10_3_17 --soloCBwhitelist BD_CLS1_v2.txt BD_CLS2_v2.txt BD_CLS3_v2.txt --runRNGseed 1 --soloMultiMappers EM --readFilesCommand zcat --outSAMattributes NH HI AS nM NM MD jM jI MC ch CB UB GX GN’ K14Cre-RBP-J floxed/floxed: ‘--soloType CB_UMI_Complex --soloAdapterSequence NNNNNNNNNGTGANNNNNNNNNGACA --soloCBposition 2_0_2_8 2_13_2_21 3_1_3_9 -- soloUMIposition 3_10_3_17 --soloBarcodeReadLength 0 --soloCellFilter None --soloCBwhitelist BD_CLS1_v2.txt BD_CLS2_v2.txt BD_CLS3_v2.txt --soloCBmatchWLtype 1MM -- soloUMIfiltering MultiGeneUMI_CR --soloUMIdedup 1MM_CR --outSAMtype BAM SortedByCoordinate --soloFeatures Gene GeneFull GeneFull_Ex50pAS --soloMultiMappers EM -- outSAMattributes NH HI AS nM NM MD jM jI MC ch CB UB GX GN --outSAMmultNmax 1 -- outMultimapperOrder Random --runRNGseed 1 --outFilterScoreMin 30 --outFilterMatchNmin 15’

#### Data Filtering, Normalisation, and Visualization

Preprocessing of the feature-count-matrix output by STARsolo was largely performed within the scanpy ^67^ecosystem, with the exception of scran normalization ^68^, which was performed in R (R Core Team, 2017). All visualization was done using ^69^ functions. Default parameters were used unless detailed otherwise.

For wild type:

Barcodes with less than 3000 UMI were discarded.

Only barcodes with less than 15% of counts stemming from mitotic genes were retained.

For K14Cre-RBP-J floxed/floxed and control:

Barcodes with less than 2000 UMI were discarded.

Only barcodes with less than 20% of counts stemming from mitotic genes were retained. Only barcodes with less than 70% of counts stemming from hemoglobin genes were retained.

And barcodes were filtered for a complexity value above 0.75, where complexity is defined as the log10-transformed number of expressed genes divided by the log10-transformed number of total counts per barcode. Scran was used for normalization.

The separate samples were merged into a single object, G2M and S phase scores were assigned to each cell using gene lists from ^70^ and the scanpy scanpy.tl.score_genes_cell_cycle function, and regressed out of the gene expression matrix using sc.pp.regress_out. 3000 highly variable genes were calculated from the normalized matrix (sc.pp.highly_variable_genes, flavor “seurat”), and the first 100 principal components were calculated with scanpy. Neighbourhood graph calculation (30 neighbors), UMAP dimensionality reduction ^71^, and Leiden clustering ^72^ were performed. scrublet ^73^ and/or scDblFinder were then used for doublet prediction.

Contaminant or undesired cell populations such as clusters of cells that, either manually or by scrublet/scDblFinder, could be classified as doublets based on their expression patterns, were removed. For this, cells were clustered at resolution 0.05, and the cluster corresponding to the epidermal cells were isolated. These were then reclustered at resolution 0.1 (wild type) and 0.2 (K14Cre-RBP-J floxed/floxed) as described above, and undesired clusters were removed. Finally, PCA, nearest neighbor calculation, and UMAP dimensionality reduction were repeated for the clean epidermis data. For WT, stages E14.5, E15.5, and E16.5 were additionally analysed separately with batch-correction using Harmony ^74^ after cell-cycle regression.

#### Differential expression analysis

Differentially expressed genes were calculated using a pseudobulk approach ^75^, comparing aggregate counts with 2 pseudoreplicates for each condition (pyDeSEQ2 0.4.8).

#### Ligand-receptor interaction analysis

We used CellphoneDB (v2.1.7 ^35^ with default parameters to analyse ligand-receptor interactions. Mouse gene names were converted to human gene names prior to the analysis.

#### Gene Set Enrichment Analysis

We ranked differential expression results by pyDESeq2 shrunken log2-fold changes and used GSEApy (v1.0.6 ^76^) “prerank” to calculate the enrichment statistics.

### Imaging Mass Cytometry

Primary antibodies from bovine serum albumin (BSA)- and azide-free formulations were conjugated using the Maxpar X8 Labeling Kit (Standard BioTools) according to the manufacturer’s protocol.

Stainings were performed on frozen cryosections according to the manufacturer’s protocol (Fluidigm). Briefly, 5 μm cryosections were prepared and stored at −80 °C until use. On the day of the experiment, slides were equilibrated at −20 °C for 1 h and then fixed in 4% paraformaldehyde (PFA) for 30 min at 4 °C. After washing in PBS, sections were incubated in blocking solution (3% BSA) for 45 min at room temperature, followed by incubation with the antibody cocktail prepared in 0.5% BSA for 1 h in a hydration chamber at room temperature. Slides were then rinsed in 0.2% Triton X-100/PBS, stained with Cell-ID Intercalator-IR (Fluidigm #201192A) for 30 min at room temperature, washed in Milli-Q water, air-dried for 30 min, and stored until imaging. The samples were laser-ablated at 200 Hz using a Hyperion imaging mass cytometry system (Fluidigm) coupled to a Helios time-of-flight mass cytometer with a resolution of 1 μm. Raw MiniCAD Design (MCD) files generated by the Hyperion Imaging System were first inspected using the MCD Viewer (Standard BioTools) to verify signal intensity and overall acquisition quality, and then exported as TIFF images for downstream analysis on Image J. ROIs delineating the basement membrane were drawn, and signal intensity at the basement membrane was measured using ImageJ software.

### Particle Imaging Velocimetry (PIV)

PIV analysis of whole-embryo live imaging was performed as previously described, with minor adjustments ^13^. Briefly, prior to analysis, each movie was processed as follows: a 3D drift correction was applied using the Fast4DReg plug-in in Fiji and sagittal cross-sections containing the basal layers were selected. A median filter of 2 pixels was applied in Fiji, followed by background subtraction using a rolling ball radius of 50 pixels.The movies were then imported into PIVLab version 2.62. Tissue flow tracking was restricted to the interfollicular epidermis. To exclude the hair follicle placodes from the analysis, ROIs covering the placodes were manually drawn on the raw images and imported into PIVLab. These masks were overlaid onto the flow field to exclude the placode regions. To prevent tracking errors due to intensity variations, image contrast was enhanced using CLAHE (Contrast Limited Adaptive Histogram Equalization) with a box size of 64 pixels.

Denoising was performed using a Wiener Noise Filter with a maximum pixel count of 3. Tissue flow tracking was carried out using a multi-pass, windowed FFT-based approach, with window sizes of 64 and 32 pixels for the first and second passes, respectively, and a 50% overlap. The resulting vector fields were visually inspected, and any erroneous vectors were manually removed from the analysis.

### Culture of primary epidermal progenitor cells

Primary epidermal cells were isolated from newborn mice essentially as described ^13^. Briefly, after dissection, skins were treated with antibiotic/antimycotic solution (Sigma A5955) for 5 min, and epidermal single cell suspensions were generated by incubating skin pieces in 0.8% trypsin for 30 min at 37 °C. Cells were suspended in Keratinocyte Growth Medium (MEM Spinner’s modification (Sigma), 5 µg/ml insulin (Sigma), 10 ng/ml EGF (Sigma), 10 µg/ml transferrin (Sigma), 10 µM phosphoethanolamine (Sigma), 10 µM ethanolamine (Sigma), 0.36 µg ml/ml hydrocortisone (Calbiochem), 2 mM glutamine (Gibco), 100 U ml/ml penicillin, 100 µg ml/ml streptomycin (Gibco), 10% chelated fetal calf serum (Gibco), 5 µM Y27632, 20 ng ml/ml mouse recombinant vascular endothelial growth factor (VEGF) and 20 ng ml/ml human recombinant fibroblast growth factor-2 (FGF-2) (all from Miltenyi Biotec) and seeded on cell culture plates coated with 30 µg ml^−1^ collagen I (Millipore) and 10 µg ml^−1^ fibronectin (Millipore) for 1 h at 37 °C.

### Micropatterned hydrogels

Circular micropatterns (100 µm diameter) were generated on glass coverslips activated by oxygen plasma treatment for 3 min (PDC-002-CE Expanded Plasma Cleaner), as described previously ^9^. Activated coverslips were incubated with 1× PLL (Surface Solutions) for 30 min at room temperature, rinsed with 10 mM HEPES (pH 7.8), and subsequently incubated with PEG-SVA (50 mg/ml in 10 mM HEPES, pH 7.8) for 1 h at room temperature. Coverslips were rinsed with Milli-Q water and air-dried before patterning by deep-UV (Primo; Alveole). PEG-cleaved regions were rinsed with Milli-Q water and incubated with 50 µg/ml fibronectin for 30 min at room temperature. In parallel, glass-bottom plates (Ibidi, #81218-200) were silanized. Polyacrylamide gels of defined stiffness were prepared by placing a drop of acrylamide/bis-acrylamide solution between the micropatterned coverslip and the silanized plate. After 45 min of polymerization, the coverslip was removed, transferring the ECM micropatterns onto the gel surface. Primary epidermal progenitors were then seeded onto the patterned hydrogels.

### Live imaging of epidermal progenitor monolayers

Freshly isolated epidermal progenitors isolated from N1IP: CreHI reporter mice crossed with R26R^mT/mG^ mice to label the plasma membrane and detect Cre activity were grown to confluency in Keratinocyte Growth Medium and plated on 35 mm glass bottom dish with 14 mm micro-well plates (Cellvis; D35-14-1.5-N). Upon confluency, 200 µM Ca^2+^ was added to induce differentiation.

Imaging was performed using an Andor Dragonfly 505 high-speed spinning disk confocal microscope (Oxford Instruments) with an inverted Nikon Eclipse Ti2 microscope (Nikon) equipped with 488 nm and 546 nm lasers, an Andor Zyla 4.2 sCMOS camera and an environmental chamber set at 37°C, 5% CO2 using a 40X dry objective. Images were acquired using Fusion 2.0 software (Andor) with multiple positions and drift stabilizer.

### Atomic Force Microscopy

For Atomic Force Microscopy (AFM), embryonic skin was dissected, immediately nitrogen frozen and then embedded in OCT blocks. AFM measurements of the basement membrane were performed on freshly cut 20 μm-thick cryosections using JPK Nano Wizard IV (Bruker Nano) atomic force microscope mounted on an Olympus IX73 inverted fluorescent microscopy (Olympus) and operated via JPK SPM Control Software v.5. Cryosections were equilibrated in PBS supplemented with protease inhibitors and measurements were performed within 20 minutes of thawing the samples.

Triangular non-conductive Silicon Nitride cantilevers (MLCT, Bruker Daltonics) with a nominal spring constant of 0,07 Nm^-1^ were used for the nanoindentation experiments. For all indentation experiments, forces of up to 3 nN were applied, and the velocities of cantilever approach and retraction were kept constant at 10 μm^s-1^ ensuring an indentation depth of 500 nm. All analysis were performed with JPK Data Processing Software (Bruker Nano). Prior to fitting the Hertz model corrected by the tip geometry to obtain Young’s Modulus (Poisson’s ratio of 0.5), the offset was removed from the baseline, contact point was identified, and the cantilever bending was subtracted from all force curves. Force curves were filtered for quality and all valid forces curves were analyzed.

### RNA in situ hybridization

RNA in situ hybridization was performed using RNAscope technology according to the manufacturer’s instructions (ACD, RNAscope Multiplex Fluorescent v2 Assay, kit: RNAscope Multiplex Fluorescent v2 Assay ACD, #323100). Briefly, paraffin-embedded tissue sections (5 μm thick) were deparaffinized in xylene, rehydrated through an ethanol series, and incubated in hydrogen peroxide for 10 min at RT. Sections were then subjected to antigen retrieval in RNAscope Target Retrieval solution for 15 min at 100°C using a pressure cooker. RNAscope Protease Plus (ACD; #322331) was applied to each section and incubated at 40°C for 30 min. The Eid1 RNAscope probe (ACD; #1713581-C3) and Keratin-DAP RNAscope probe (ACD; #500671-C1) were hybridized according to the manufacturer’s protocol using the RNAscope Multiplex Fluorescent v2 Assay (ACD; #323100), followed by signal detection with TSA Plus Fluorescein (Akoya, #NEL741001KT), TSA Plus Cyanine 3 System (Akoya, #NEL744001KT). All hybridization and amplification steps were performed in a HybEZ II oven (ACD). DAPI solution was incubated for 30 seconds at room temperature. Sections were mounted in Elvanol and imaged using a Zeiss LSM 980 confocal microscope equipped with a 40× water-immersion objective, using Zeiss ZEN software (version 3.5).

### Statistics and reproducibility

Statistical analyses were performed using GraphPad Prism software (GraphPad, version 9) or in R (version 4.2.2). Statistical significance was determined by the specific tests indicated in the corresponding figure legends. Only 2-tailed tests were used. In all cases where a test for normally distributed data was used, normal distribution was confirmed with the Kolmogorov–Smirnov test (α = 0.05). All experiments presented in the manuscript were repeated at least in 3 independent replicates.

## Data availability

Sequencing datasets are available at GEO (GSE315748 and GSE315746). All analysis scripts, custom code, and data that support the conclusions are available from the authors on request.

## Acknowledgements

We thank Claudia Ortmeier, Manuela Haustein, Hermann vom Bruch, and Sandra Heising for expert technical assistance, Silvia Fre (Institute Curie, France) for advice on the project and general support, Fabien Bertillot for advice on image analyses, Yekaterina Miroshnikova for advice on micropatterning, and the Max Planck Institute BioOptics and Sequencing core facilities for support with experiments. Imaging Mass Cytometry was performed at the Cell Imaging and Cytometry Core, Turku Bioscience Centre (Turku, Finland), with the support of Biocenter Finland. This work was supported by European Union’s Horizon 2020 research and innovation programme under Marie Skłodowska-Curie grant agreement no. 101032331 (to CV), Research Council of Finland (332402) and The Turku Collegium for Science, Medicine, and Technology (to GF), Academy of Finland Center of Excellence BarrierForce (346131) and R’Life Programme consortium NucleoMech (330033) (to JI and SAW) the Sigrid Juselius Foundation, Helsinki Institute of Life Science, and the Max Planck Society (all to SAW), NSF-CMMI-1334611 (to SA and MLM), grant number 2023- 329572 from the Chan Zuckerberg Initiative DAF, an advised fund of the Silicon Valley Community Foundation (to MLM), and the Syracuse University Office of Research Future Faculty Fellow Program (to SA). CMN is supported by the Deutsche Forschungsgemeinschaft (DFG, German Research Foundation: Germanýs Excellence Strategy – EXC 2030 – 390661388; SPP 1782 NI 1234/6-2, Project-ID 388932620; FOR 2743 NI 1234/7-1 Project-ID 678823; ANR/DFG NI 1234/9 (Project-ID 505673300) and the European Union NETSKINMODELS, COST Action no. CA21108.

## Author contributions

CV, MA, and MR designed and performed experiments and analyzed data. SHAE and MLM developed vertex model and performed all simulations and analyses. KK analyzed sequencing data. SV and LCB supported live imaging and single cell sequencing experiments. AP performed and analyzed immunofluorescence experiments, PZ assisted with imaging. GF and JI performed mass cytometry experiment. CMN, MLM and SAW conceived and supervised the study, designed experiments, and analyzed data. MLM and SAW wrote the paper. All authors commented and edited the manuscript.

## Supplementary Figures

**Supplementary Figure 1.**
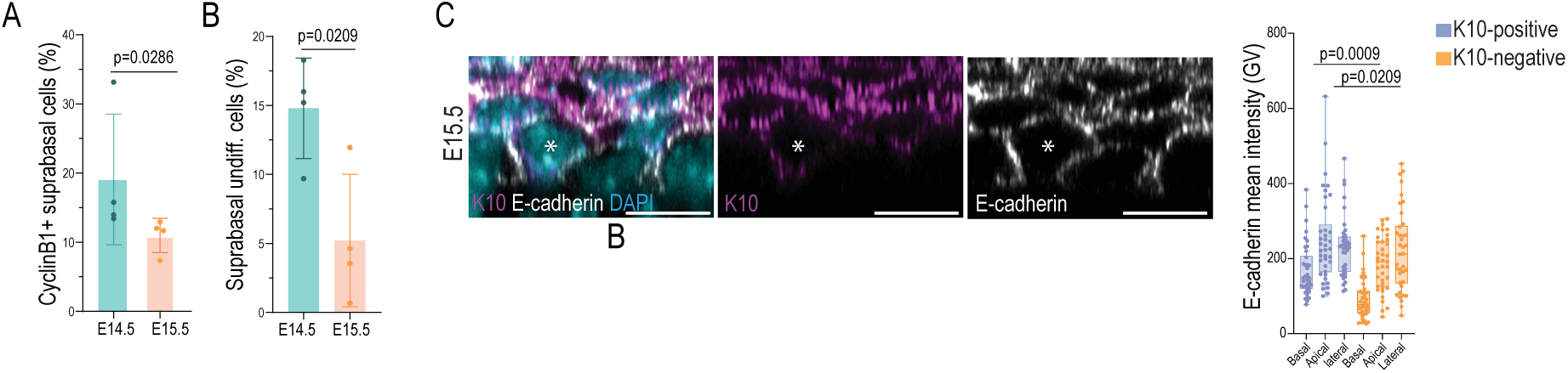
**A.** Quantification of differentiated (K10-positive) CyclinB1+ cells in the suprabasal layer (mean ±SD; n=4 embryos /stage; Mann-Whitney) **B.** Representative images and quantification of delaminating cells in E15.5 embryo from 3D whole mounts stained against E-cadherin, K10 and DAPI. Delaminating cell is marked with a white asterisk; scale bars 10 µm; min-max box and whiskers; n=38 cells pooled across 3 embryos; Friedman/Dunn’s).

**Supplementary Figure 2.**
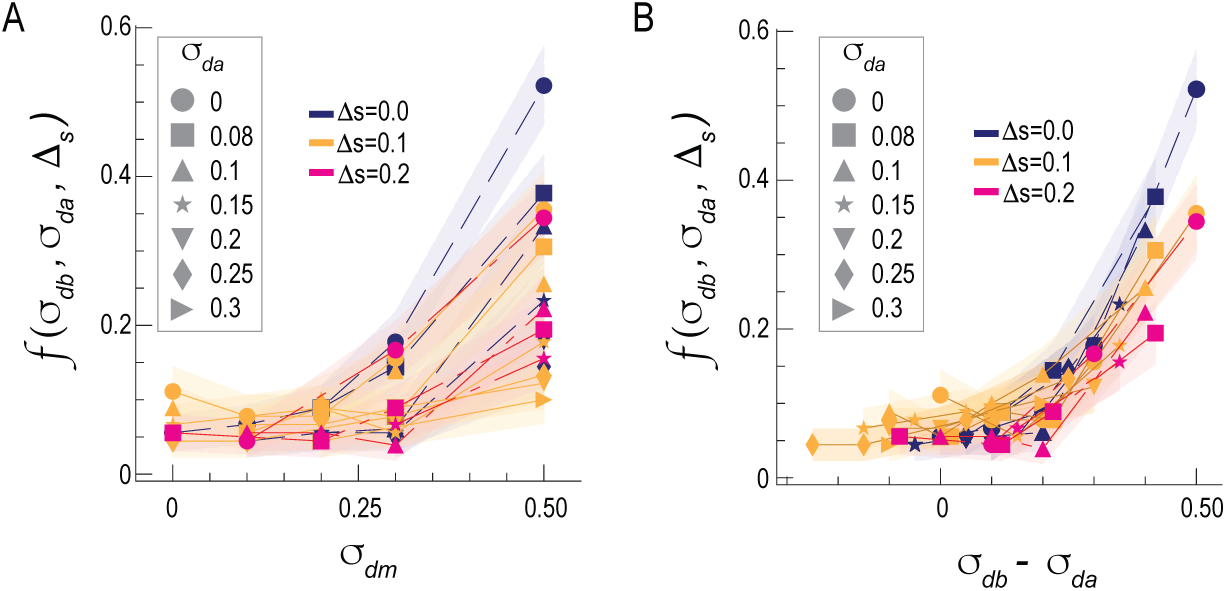
Systematic quantification of delamination rates as a function of the properties of the delaminating and stiffness of the surrounding tissue 𝚫𝒔. **A.** Uncollapsed delamination rates. **B.** Delamination rates plotted as a function of 𝜎_𝑑𝑏_ − 𝜎_𝑑𝑎_, which do not fully collapse, indicating that an additional energetic contribution is required. Figure 2D in the main text shows that adding the term 𝑐 Δ𝑠leads to a collapse of the curves onto a single master curve as a function of 𝜎_𝑑𝑏_ − 𝜎_𝑑𝑎_ + 𝑐 Δ𝑠.

**Supplementary Figure 3.**
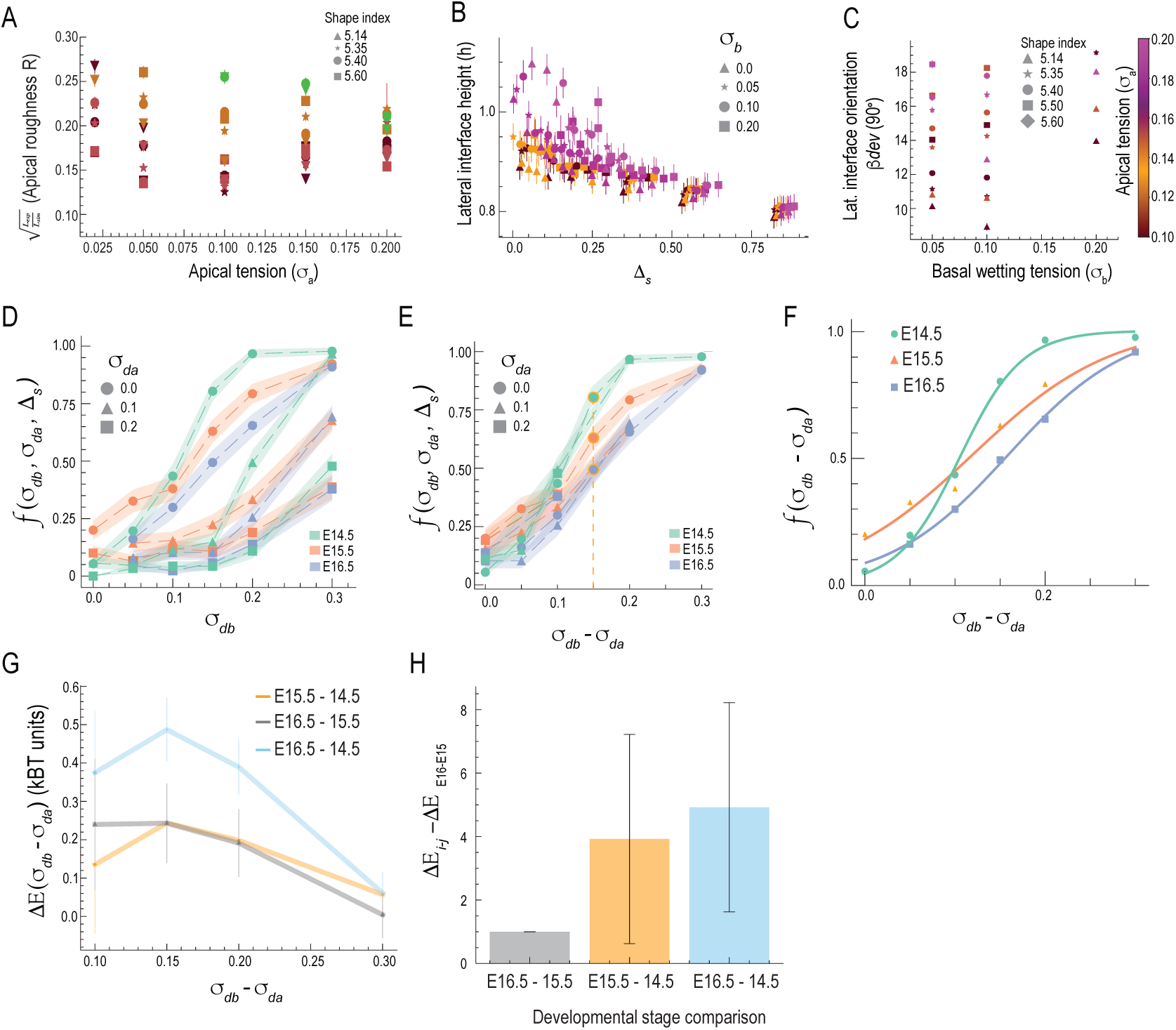
Geometric observables for mechanical barrier and quantification of delamination rates and associated energy barriers as function of delaminating cell properties. **A-C.** Uncollapsed plots of basal-suprabasal interface roughness, basal cell lateral interface height, and lateral interface orientation, respectively, measured in the 3D model simulations across mechanical parameter space (𝜎_𝑎_, 𝜎_𝑏_, Δ𝑠). Corresponding collapsed plots are shown in main Fig. 3G, I, K. **D.** Quantification of delamination rates data as function of the delaminating cell tensions (𝜎_𝑑𝑏_, 𝜎_𝑑𝑎_) parameter space. **E.** Data from (F) collapses as a function of 𝜎_𝑑𝑏_ − 𝜎_𝑑𝑎_. **F.** Curve fits to the collapsed delamination rate data shown in panel (B) across the developmental stages. **G.** Quantification of the differences in energy barriers between developmental stages as a function of the differences in delaminating cell properties (𝜎_𝑑𝑎_, 𝜎_𝑑𝑏_). **H.** Normalized averages (normalized by the energy barrier difference between stages E16 and E15) of the differences in energy barriers between stages E15.5 and E14.5, and between E16.5 and E14.5, plotted as a function of 𝜎_𝑑𝑏_ − 𝜎_𝑑𝑎_.

**Supplementary Figure 4.**
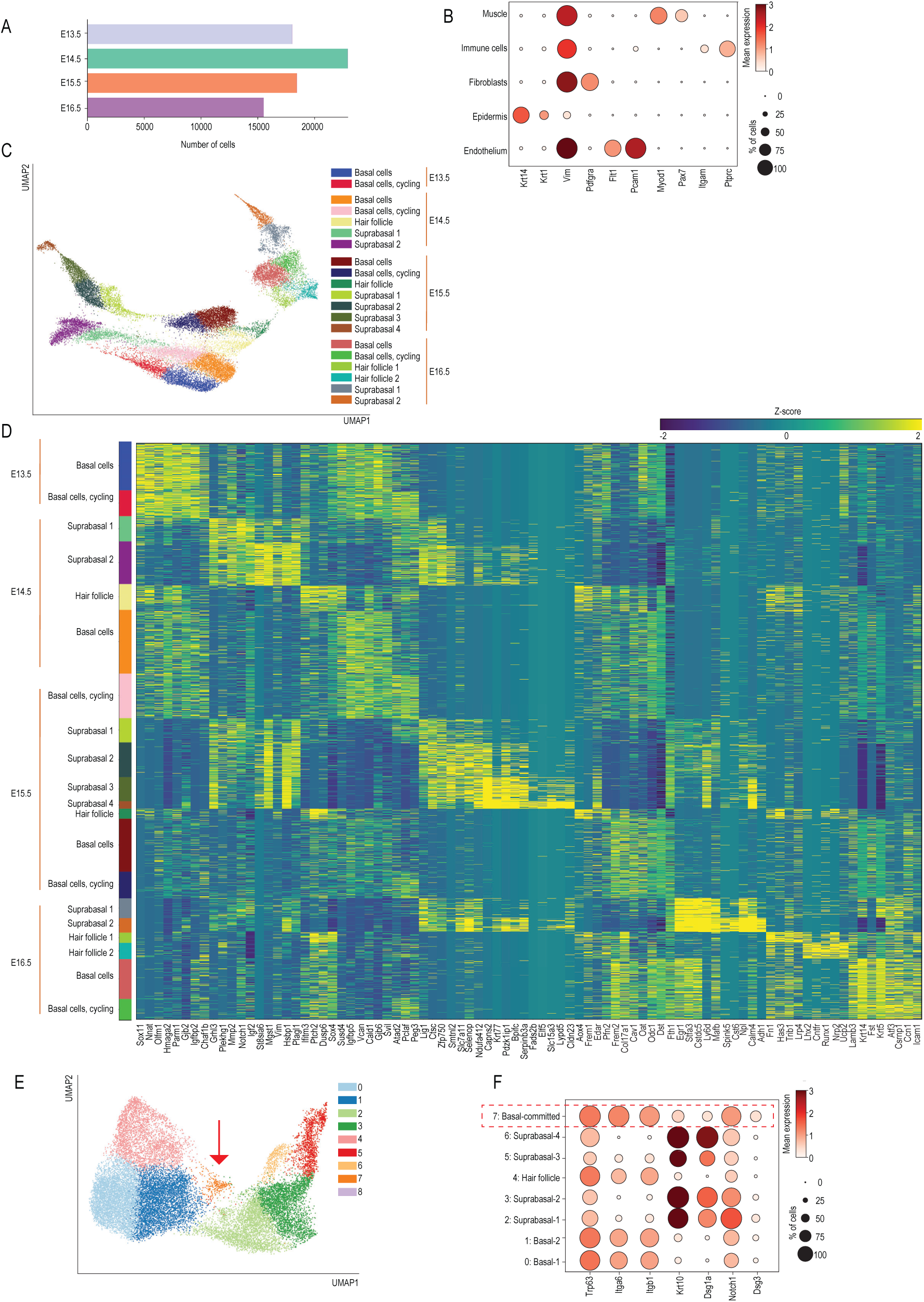
**A.** Bar plot showing number of sequenced cells/genotype. **B.** Dot plot of selected key cell type-specific marker genes across annotated cell types. **C.** UMAP visualization of Leiden clusters of epidermal cells with main cell states annotated. **D.** Heatmap of marker genes for epidermal Leiden clusters. **E.** UMAP visualization of Harmony integrated epidermal cells. Red arrow indicates committed cell population. **F.** Dot plot of selected key cell type-specific marker genes across annotated cell types from Harmony integrated epidermal cells. Red dotted rectangle indicates committed cell population.

**Supplementary Figure 5.**
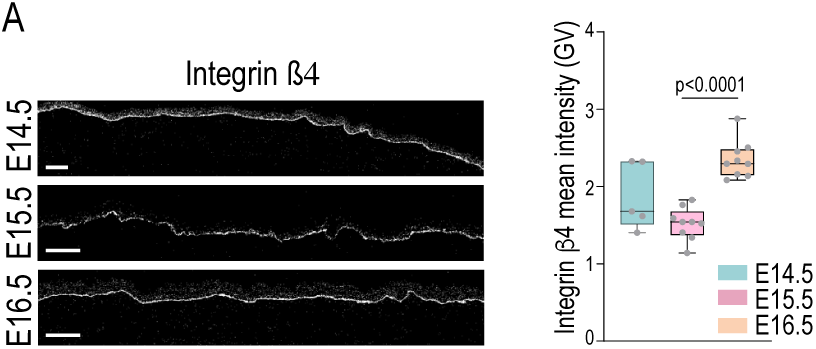
**A.** Representative images and quantifications of tissue sections from E14.5- 16.5 embryos analyzed by mass cytometry. Note increased expression of integrin β4 (mean ±SD; scale bars 50 µm; n=3 embryos/stage; Two-way ANOVA/Tukey’s.

**Supplementary Figure 6.**
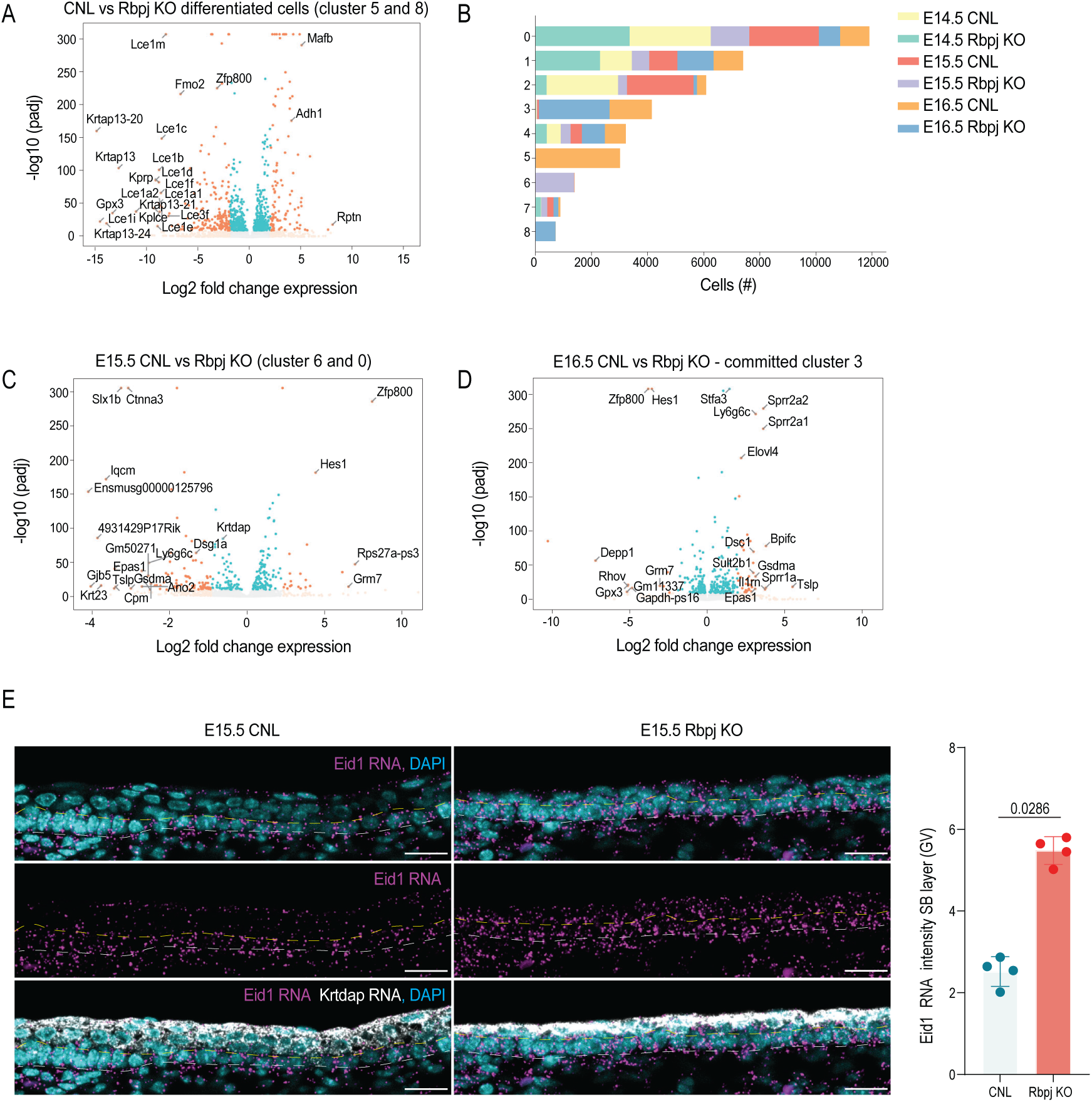
**A.** Volcano plot showing differentially expressed genes between Rbpj-KO and CNL epidermis from differentiated cell clusters 5 and 8. **B.** Bar plot showing number of sequenced cells/embryonic stage and genotype across Leiden clusters. **C.** Volcano plot showing differentially expressed genes between E15.5 Rbpj-KO and CNL epidermis from clusters 6 and 0. **D.** Volcano plot showing differentially expressed genes between E16.5 Rbpj-KO and CNL epidermis from cluster 3. **E.** Representative images and quantification of tissue sections from E15.5 embryos of control and Rbpj-KO mice of the expression of Eid1 (magenta) and Keratin-dap (grey) detected by RNA in situ hybridization. Dotted lines mark basal layer (n=4 embryos/condition; Mann-Whitney). Scale bars 20 µm.

